# Chromatin organization by an interplay of loop extrusion and compartmental segregation

**DOI:** 10.1101/196261

**Authors:** Johannes Nuebler, Geoffrey Fudenberg, Maxim Imakaev, Nezar Abdennur, Leonid Mirny

## Abstract

Mammalian chromatin is organized on length scales ranging from individual nucleosomes to chromosomal territories. At intermediate scales two dominant features emerge in interphase: (i) alternating regions (<5Mb) of active and inactive chromatin that spatially segregate into different compartments, and (ii) domains (<1Mb), i.e. regions that preferentially interact internally, which are also termed topologically associating domains (TADs) and are central to gene regulation. There is growing evidence that TADs are formed by active extrusion of chromatin loops by cohesin, whereas compartments are established by a phase separation process according to local chromatin states. Here we use polymer simulations to examine how the two processes, loop extrusion and compartmental segregation, work collectively and potentially interfere in shaping global chromosome organization. Our integrated model faithfully reproduces Hi-C data from previously puzzling experimental observations, where targeting of the TAD-forming machinery led to changes in compartmentalization. Specifically, depletion of chromatin-associated cohesin reduced TADs and revealed hidden, finer compartments, while increased processivity of cohesin led to stronger TADs and reduced compartmentalization, and depletion of the TAD boundary protein, CTCF, weakened TADs while leaving compartments unaffected. We reveal that these experimental perturbations are special cases of a general polymer phenomenon of active mixing by loop extrusion. This also predicts that interference with chromatin epigenetic states or nuclear volume would affect compartments but not TADs. Our results suggest that chromatin organization on the megabase scale emerges from competition of non-equilibrium active loop extrusion and epigenetically defined compartment structure.

## Introduction

Eukaryotic chromatin, i.e. DNA together with associated proteins, is far from being simply a randomly arranged polymer in the cell nucleus. Investigations into its spatial organization by chromosome conformation capture (1) and its descendent Hi-C (2) have revealed two salient features in higher eukaryotes. First, at the super-megabase (Mb) scale, chromatin spatially segregates into different compartments (2). The Hi-C signature of segregation is a plaid, or checkerboard pattern (Fig. 1A), which indicates that chromatin of a given type preferentially interacts with other loci of the same type (3, 4). Spatial segregation is further supported by imaging of individual loci (5, 6) and whole compartmental segments (7). The second striking feature of 3D organization are topologically associating domains (TADs) (8, 9). Their Hi-C signature are squares along the diagonal, indicating local regions of increased contact frequency, typically on the sub-Mb scale.

Several lines of evidence indicate that compartments and TADs are formed by distinct mechanisms and are not a hierarchy of the same phenomenon on different scales. First, TADs have no checkerboard pattern in Hi-C (Fig. 1, and (8)). Second, the alternating compartment structure correlates with gene density, gene expression and activating epigenetic marks, which are all enriched in compartments of type A (2), while no such classification has been reported for TADs. Rather, TAD boundaries, not their interior, are associated with architectural proteins, in particular CTCF (8, 9). Also, TADs are less cell type-specific than compartments (8, 9). Furthermore, TADs can exist without compartments and vice versa (10). And finally, recent experiments directly showed that TADs compete with compartments: removal or depletion of chromatin-associated cohesin (11–14), which is required for TADs, not only made TADs disappear but also increased compartmentalization (11, 13, 14), sharpened compartment transitions (12), and fragmented compartments into shorter intervals (11) (see Fig. 1B for a cartoon and Fig. 2A for an example). Strikingly, these finer compartments match epigenetic marks of activity better than the more coarse wild-type compartments (11), suggesting that the loss of cohesin activity reveals underlying innate compartment structure that is obscured in the wild type (WT). The opposite effect was achieved by increasing the residence time and the amount of cohesins on DNA: TADs were extended and compartmentalization weakened (13, 14) (see Fig. 4A for an example). These observations raise the question of how cohesin, crucial for forming TADs, could mechanistically alter compartmentalization.

TAD are believed to be formed by active extrusion of chromatin loops (15, 16), which has appeared multiple times in the literature as a mechanism for chromosome organization (17–20): loop extrusion factors (LEFs) attach to the chromatin fiber and start progressively enlarging a DNA loop until they either fall off, bump into each other, or bump into “roadblocks”, which define the TAD boundaries (Fig. 1B). Active loop extrusion explains many features of TADs (15, 16): (i) TADs have no checkerboard signature in Hi-C, (ii) removal of a TAD boundary leads to the fusion of two TADs into a larger one, (iii) the sequence motifs at TAD boundaries have a specific, convergent, orientation as they oppose loop extrusion unidirectionally, and (iv) TAD corner peaks arise from loop extruders bringing TAD boundaries into spatial proximity.

**Fig. 1.**
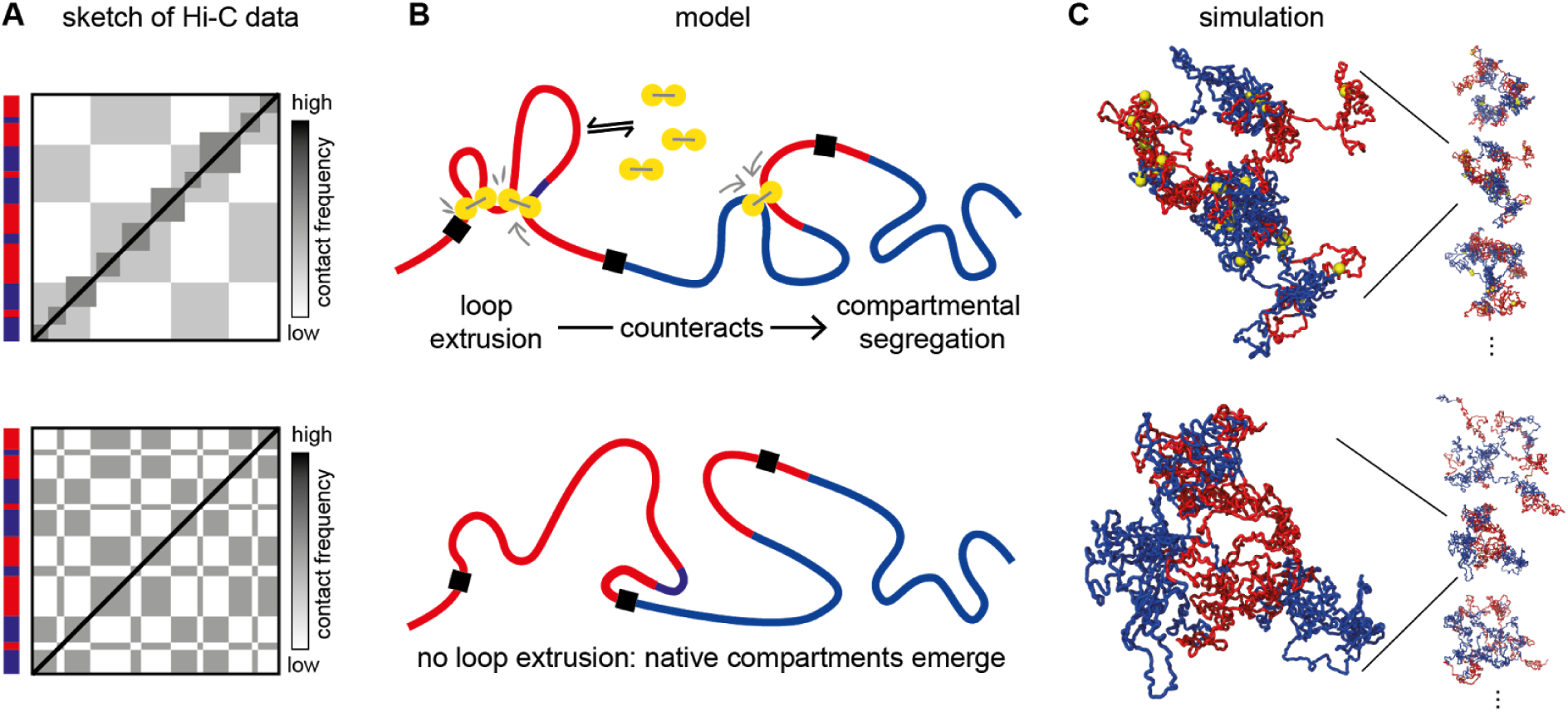
Loop extrusion competes with compartmental segregation. (A) Cartoon of typical Hi-C signatures: topologically associating domains (TADs) are squares of increased contact frequency along the diagonal, while compartmentalization is as a checkerboard pattern indicating spatial segregation. Upon removal of the cohesin loader Nipbl, Schwarzer et al. observed that TADs disappear and a fine scale compartmentalization emerges (indicated in red/blue on the left, see Fig. 2A for a data example). The decay of the average contact probability with linear genomic distance (scaling) is not shown in the cartoons. (B) Sketch of our mechanistic model: loop extrusion factors (LEFs, yellow) counteract compartmental segregation. (C) Example conformations from polymer simulation ensembles (shown are 10 Mb sections from our simulated 50 Mb fibers).

The proposed molecular candidates for LEFs are Structural Maintenance of Chromosome (SMC) protein complexes (21, 22), in particular cohesin during interphase (23). Cohesin topologically entraps DNA (24), can slide along DNA and over small DNA bound proteins and nucleosomes (25, 26) and is enriched at TAD boundaries (9) and corner peaks (27). Recently, Terakawa et al. showed that a closely related SMC, namely yeast condensin, processively walks along DNA consuming ATP *in vitro (28)*. Furthermore, bacterial condensins processively juxtapose the bacterial chromosome *in vivo* (29). Cohesin is loaded onto eukaryotic DNA, assisted by Nipbl (30), while WAPL limits its residence time (31, 32). Central to the formation of TAD boundaries is the protein CTCF: it is enriched and conserved at TAD boundaries (8, 9) and disruption of CTCF binding sites alters TAD structure (15, 33–36).

Compartmental segregation, suggested by a checkerboard pattern in Hi-C maps, can not be explained by loop extrusion. Rather, a natural class of models are block-copolymers (37, 38): polymers consisting of alternating regions which differ in contact interaction exhibit segregation of different blocks into separate spatial compartments (39–41). These models are further motivated by the observed partitioning of chromatin into a small number of types based on DNA binding protein profiles and modifications (27, 42), which may in turn entail differences in contact interaction, including via histone tails (43) and recruitment of HP1 proteins (44, 45). An integrated model that includes both compartmentalization and loop extrusion is largely missing. While Rao et. al. (2017) illustrate how the pattern of compartments and TADs change in simulations upon loss of loop extrusion in a single 2Mb locus, a systematic characterization and a physical examination of how the non-equilibrium active loop extrusion process affects global compartmentalization is essential for understanding large-scale chromosome organization.

Here we address the question of how cohesin-mediated loop extrusion can affect compartmentalization of hetero- and euchromatin, in other words how an active process can interfere with the phase separation of a block copolymer. Using polymer simulations we show that active mixing by loop extrusion locally counteracts the phase separation. Our model agrees with several recent experiments where targeting of different parts of TAD-forming system had different effects not only of TADs but also on compartments. Our model also makes specific predictions for future experiments and explains how the interplay of loop extrusion and compartmental segregation shapes chromosome organization in interphase.

## Results

### Polymer model of loop extrusion and compartmental segregation

In order to investigate the interplay of loop extrusion with compartmentalization we simulate the chromatin fiber as a polymer subject to loop extrusion and compartmental segregation (Fig. 1A). Loop extrusion factors (LEFs) can attach to the chromatin polymer at random positions and extrude loops bidirectionally until they either fall off, bump into each other or encounter a boundary element. When blocked on one side they continue extruding unidirectionally. LEFs are characterized by three parameters: the average residence time τ the single-sided extrusion velocity *v*, and the average separation *d* (46)*. The first two define the processivity as λ =2τv*, which is the average size of a loop extruded by an unobstructed LEF. For our simulation of wild type (WT) cells we use a processivity of λ =250 kb and a separation of *d*=750 kb. As λ /*d*=1/3 our WT cells operate in the dilute regime (λ /*d*<1) where LEFs rarely bump into each other. CTCF-enriched boundaries of TADs are modeled by elements that block extrusion of LEFs with probability 90% (16). Having a finite permeability is consistent with the turnover time of CTCF being considerably shorter than that of cohesin (~1-2min (47, 48) vs. >30min (14, 48–51)), though the exact value of the permeability is yet to be determined and may depend on the number and occupancy of CTCF sites at each boundary and molecular details of interactions between CTCF and cohesin. Values of these and other parameters are chosen to reproduce TAD patterns observed in Hi-C data and systematically varied to examine their effects on chromatin organization.

Compartment organization is modeled by a block-copolymer composed of A and B blocks that have the same local properties (monomer size and fiber flexibility) but interact differently. The spatial segregation is induced by weak B-B attraction (referred to as *compartmental interaction*, see Fig. S1), but can also be modeled differently (Fig. S2). To reflect the fine compartment structure revealed by depletion of chromatin-associated cohesin (11), we assigned alternating compartmental segments that are typically larger than TADs, but interspersed with small regions of the respective other type (see Supplement). Lengths of A and B regions were chosen to yield an autocorrelation length of the compartment profile that is consistent between experiments and simulation (see Supplement). The TAD and compartmental layouts in simulations are randomly generated based on the above characteristics and are not intended to reproduce specific genomic regions.

To clarify compartment related terminology: in simulations, each locus is either of *type* A or B (see above), depending on its interaction preference with other loci. We refer to a checkerboard pattern in Hi-C maps as *compartmentalization* and to the corresponding colocalization in 3D as *compartmental segregation*. Furthermore, we compute *compartment profiles* from eigenvector decomposition of a Hi-C matrix (2, 3). *Compartmental segments* of type A/B are intervals where the eigenvector is positive/negative. Note that a locus of a given type may not be able colocalize with other loci of the same type. Compartmental segments may thus differ from the underlying types of loci. We measure the *degree of compartmentalization* of a Hi-C map as the normalized excess of contacts between loci of the same type over contacts between loci of different types, (AA+BB-AB)/(AA+BB+AB), (see Supplement). With experimental data, the compartment profile is used to assign A/B status to loci since the underlying types are not known. For consistency, we do the same for simulated Hi-C maps when measuring the degree of compartmentalization.

Unless otherwise mentioned, we allow for some passing of two parts of the chromatin fiber through each other by imposing a finite repulsive core on the monomer interaction potential (Fig. S1). This represents the effect of topoisomerase II and is discussed further below.

### Loop extrusion overrides compartmentalization on small scales

We first test whether our integrated model of loop extrusion and compartmental segregation can explain the effect of depletion of chromatin-associated cohesin (11), namely disappearance of TADs and simultaneous changes in compartmentalization such as (i) compartmental segments that span several Mb to several tens of Mb appear more crisp in Hi-C, and (ii) they become fragmented into smaller segments (Fig. 2A). Strikingly, loss of loop extrusion in our model reproduces both phenomena: while TADs disappear, compartmentalization, in particular of small segments, is enhanced, leading to fragmentation of large compartmental segments (Fig. 2B). Our simulations thus show that loop extrusion suppresses the inherent compartmentalization by counteracting segregation of small segments, which emerges when loop extrusion is removed.

We quantify changes in simulated chromatin upon loss of loop extrusion and compare them to changes in experimental data from (11) in three ways (Fig. 2A and B, bottom panels). (i) The removal of loop extrusion is detected by changes in the contact frequency as a function of genomic distance, P(s). With loop extrusion the *P(s)* curve shows a characteristic hump on the length scale of TADs. This hump disappears upon removal of loop extrusion both in experiment and simulations. (ii) The strengthening of short compartmental segments (“fragmentation” of compartments) upon loss of loop extrusion is quantified by the steeper decay of the autocorrelation of the compartment profile. This steepening is evident in simulations and experiment alike. (iii) The greater contrast in Hi-C maps upon removal of loop extrusion is measured by changes in the degree of compartmentalization (see above and Supplement). Its increase in simulations is slightly stronger than in experiments, which could indicate that some compartment mixing remains present in experiments, either by residual cohesin (see Fig. S4) or some other processes in the nucleus not considered here (Supplemental Note).

Most importantly, our simulations show that loop extrusion suppresses small compartmental segments more than large ones (see Fig. 2C). This explains how cohesin depletion experiments reveal the fine scale compartmentalization: larger compartmental segments remain unchanged, while small ones that are suppressed by loop extrusion in WT cells emerge in the mutant without loop extrusion. Our simulations suggest that the emergent fine structure is the intrinsic compartmentalization that is overridden in WT cells by cohesin activity. This is in line with the observation that epigenetic marks correlate better with finer emergent than with the coarser WT compartmentalization (11). As a specific example, we see how a small compartmental segment that spans across a TAD boundary can emerge upon removal of loop extrusion both in experiment and simulation (Fig. 2D).

Taken together, our results suggest that loop extrusion suppresses the inherent compartmental segregation on the length-scale of loops and leaves only larger scale compartmentalization visible. When loop extrusion is removed by depletion of chromatin-associated cohesin, the intrinsic compartmental segregation emerges.

**Fig. 2.**
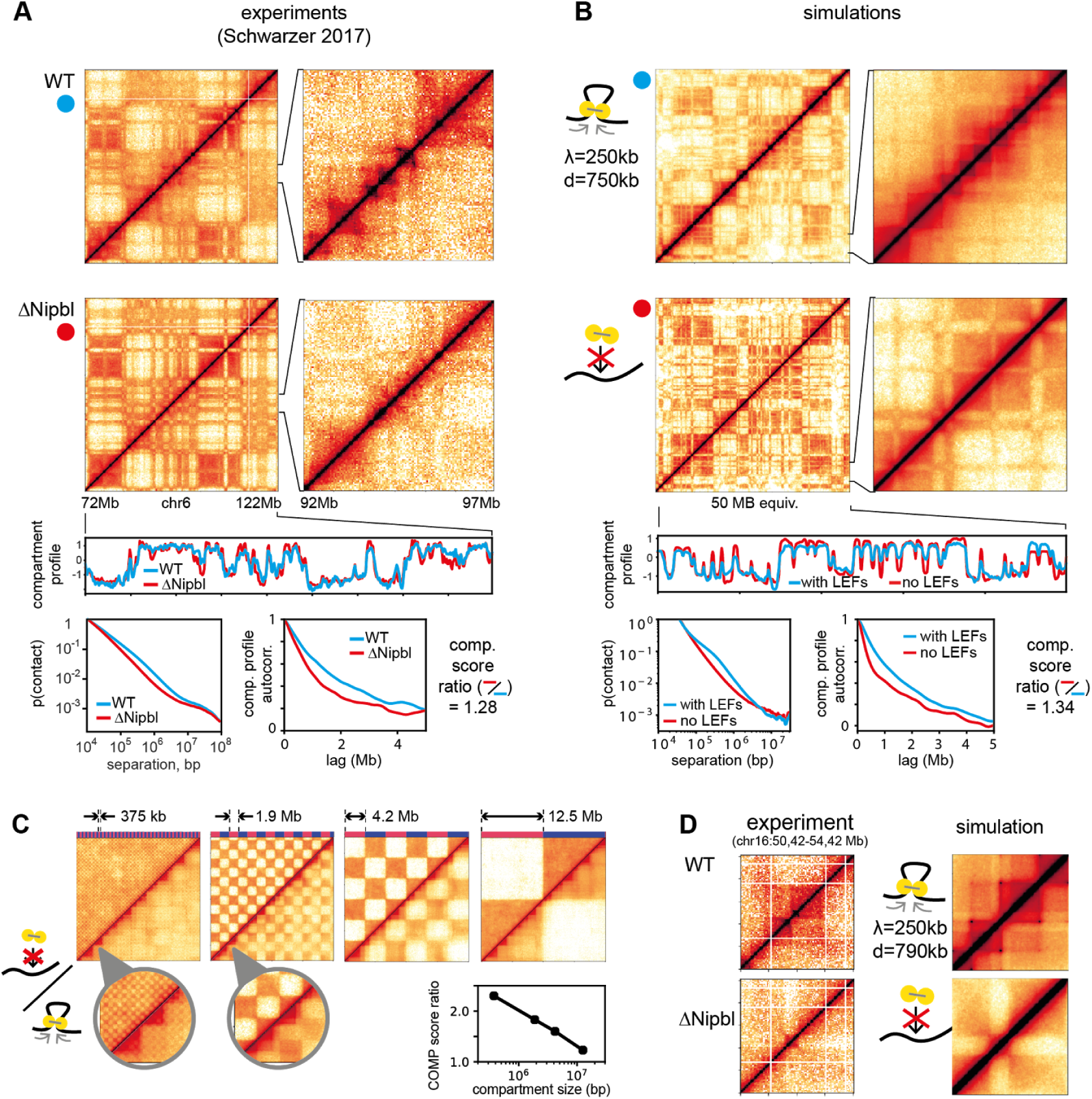
Removal of Nipbl -- removal of loop extrusion. (A) Removal of DNA associated cohesin by deletion of Nipbl leads to stronger and fragmented compartmentalization and loss of TADs. (B): The same is observed in simulated Hi-C maps upon removal of loop extrusion. The loss of loop extrusion leads the loss of a characteristic hump in the contact probability as a function of genomic separation. The fragmentation is apparent in compartment profiles as the faster decay in their autocorrelation. The degree of compartmentalization (comp score) is reduced by a similar factor upon removal of Nipbl/loop extrusion in experiments/simulation. The scaling for experimental Hi-C maps is computed genome wide, other quantities on the shown regions. The TAD and compartment arrangement is randomly generated and not intended to reproduce the experimental example. (C) Large compartmental segments are affected less by loop extrusion than small ones: upper/lower triangles: with/without loop extrusion. (D) Upon removal of loop extrusion, a previously hidden compartmental segment emerges that spans two TADs, both in experiment and simulations. All panels: Throughout the text contact frequency is shown on a log-scale with bounds adjusted for optimal feature visibility. Resolutions: 200kb for 50Mb maps, 40kb for 5Mb maps, 100kb for compartment profiles.

### Removing loop extrusion boundaries suppresses TADs but not local compaction or compartments

Next, we asked if our model of loop extrusion and compartmental segregation is compatible with depletion experiments of the TAD boundary element CTCF (36). Namely, CTCF depletion leads to a loss of TAD boundaries while having little effect on compartmentalization (Fig. 3A). We simulated CTCF depletion by removing boundary elements (Fig. 3B), which led to a 1.2-fold increase in loop size (from 173 kb to 216 kb). In agreement with experiments we observe a loss of TADs, while compartmentalization is affected only slightly (Fig. 3B, see Fig. S5 for a parameter sweep). Unlike in cohesin depletion, no fine compartmentalization emerges. The distinction from cohesin depletion arises because upon CTCF removal loop extrusion is still present, but not restricted to specific domains.

Although TADs are diminished upon both CTCF removal and cohesin loss, these two perturbations have vastly different effects on chromatin organization. The lack of changes in *P(s)* curves upon CTCF removal suggest that loop extrusion is unaffected. It remains locally compacted, as evident from the hump for s<1Mb in the P(s) curve. Our simulations reproduce this phenomenon: the loss of boundaries while maintaining loop-extrusion removes TADs but preserves *P(s)*. Loss of chromatin-associated cohesin in experiment, on the contrary, leads to the reduced compaction as evident by the loss of the hump in the P(s) curve. Simulations with diminished loop extrusion activity reproduced these changes (see above and Fig. 2). Corresponding changes in compartmentalization upon cohesin loss and the lack of such changes upon CTCF removal suggest that it is the loop extrusion activity of cohesin that lead to coarsening of compartmentalization in the wild-type.

**Fig. 3.**
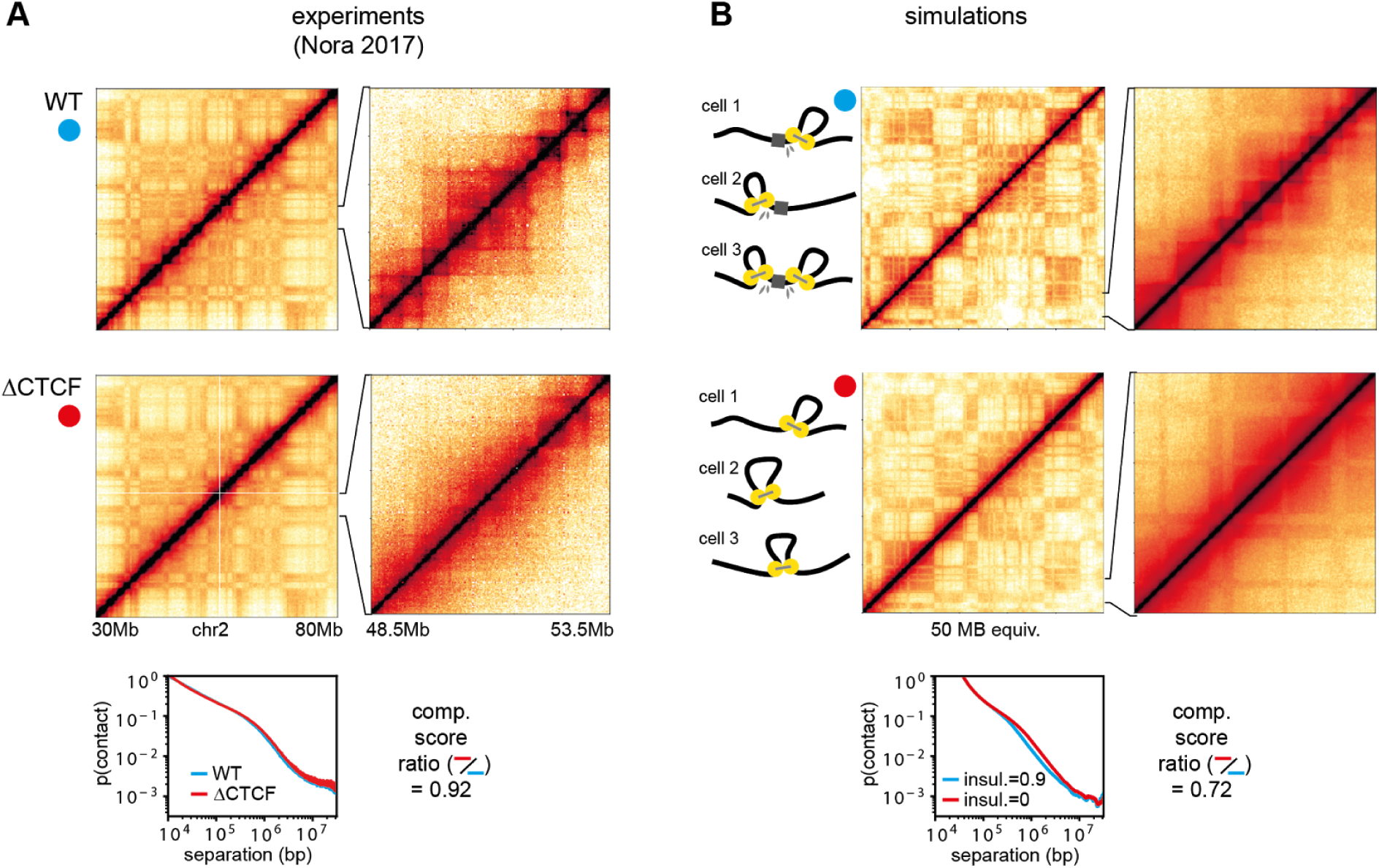
Removal of CTCF -- removal of boundary elements. (A) CTCF depletion strongly suppresses TADs but leaves compartmentalization almost unaffected (data from (36)). (B) The same is observed in simulations when loop extrusion boundaries are removed (boundary insulation reduced from 90% to 0%). Processivity λ and separation *d* are as in Fig. dNIPBL. The decay of the contact probability with genomic distance barely changes both in experiment and simulation. The degree of compartmentalization is reduced slightly upon removal of CTCF/boundaries.

### Increasing loop extrusion processivity suppresses compartments while enhancing TADs and secondary corner peaks

Finally, we consider how increased processivity of cohesin can affect compartmentalization and examine the experimental removal of the cohesin release factor Wapl. Wapl removal weakens compartmentalization and strengthens TADs and corner peaks (Fig. 4A) (13, 14). It was reported that the amount of chromatin-associated cohesin in Wapl deficient cells increases moderately (~1.5-2-fold), while the residence time increased considerably (>5-fold) (13, 14). We thus simulated Wapl removal by increasing the LEF density 1.5-fold (reducing the average separation from 750 kb to 500 kb) and the residence time ten-fold, which results in larger processivity (2.5 Mb instead of 250 kb). The average loop size increased only 2.6-fold (from 173 kb to 449 kb), indicating that extrusion becomes limited by LEFs bumping into each other (as sketched in Fig. 4B). In agreement with experiments, this leads to stronger TADs and more pronounced corner peaks (Fig. 4B). Corner peaks between non-adjacent boundaries are particularly enhanced (Fig. 4B, lower sketch). The change in the contact probability *P(s)* in our simulations is also consistent with changes in experimental *P(s)* curves (Fig. dWAPL, bottom panels) which show an extension of the characteristic hump to larger genomic separations, reflecting larger extruded loops.

Also in agreement with experiments, our simulations of Wapl removal show reduced compartmentalization (Fig. 4B left panels). We attribute this to increased compartment mixing by the longer and more abundant loops. Further suppression of compartments in Wapl-depleted cells might be due to formation of axially compressed and stiff “vermicelli” chromosomes (50), which can limit far-cis contacts and interactions with the lamina, thus affecting compartmentalization.

**Fig. 4.**
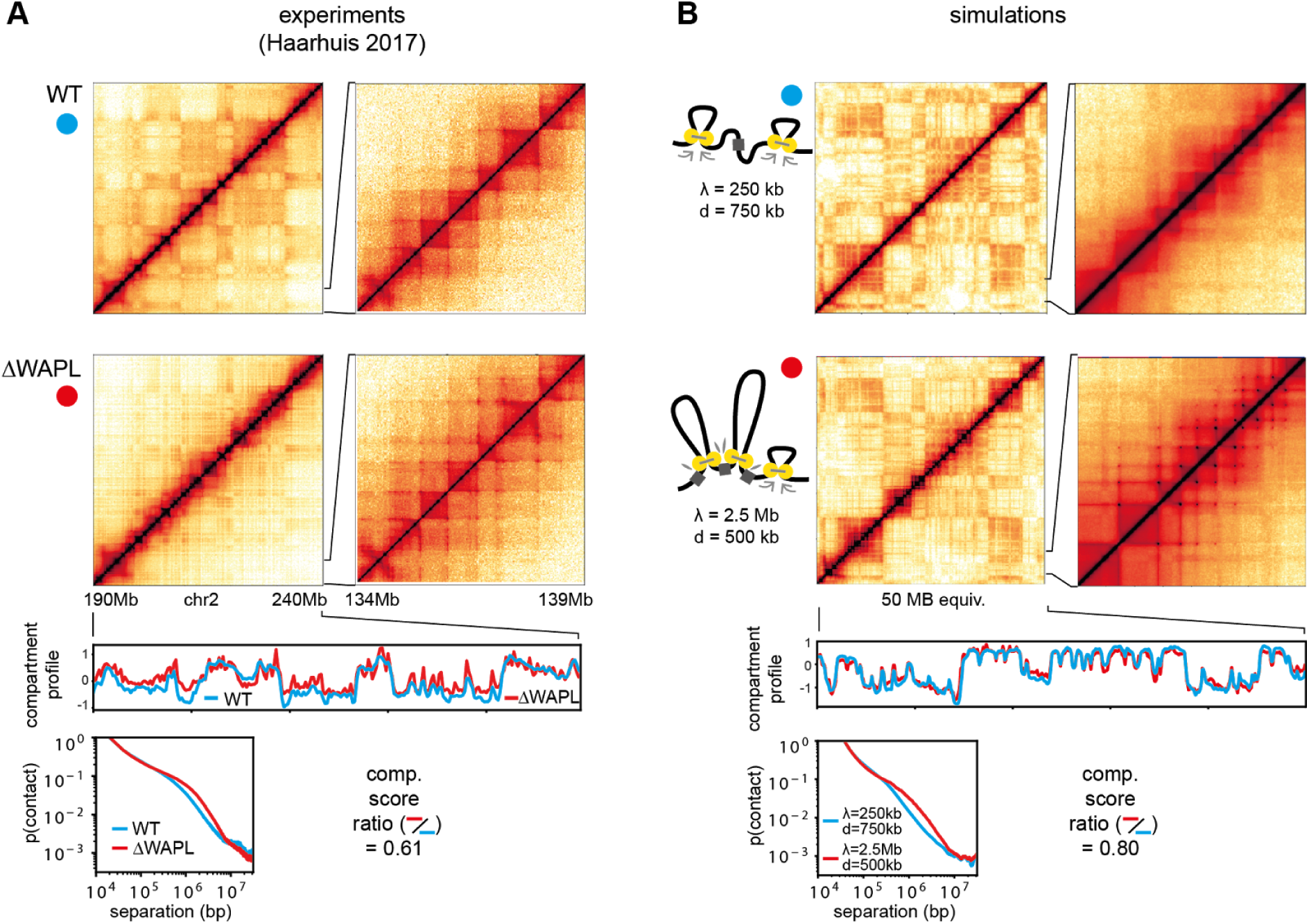
Removal of Wapl -- enlarged loops. (A) Removal of Wapl reduces compartments and strengthens TADs, in particular secondary corner peaks. Data from (14). (B) The same is observed in simulations with a 10 fold increase in LEF residence time and a 1.5 fold increase in LEF density. The secondary corner peaks arise from fully extruded TADs, forming contacts between several consecutive boundary elements (lower cartoon). The characteristic hump in contact probability scaling extends to significantly larger loops. The degree of compartmentalization is reduced.

### The non-equilibrium nature of compartment mixing by loop extrusion

We have shown above that compartment mixing by loop extrusion explains the changes of TADs and compartmentalization for all considered experimental perturbations. We thus aim at understanding physical mechanisms behind this mixing effect.

Loop extrusion brings loci into contact irrespective of their compartmental identity. We thus asked if the reduced compartmentalization due to loop extrusion can be simply understood by an effective reduction of the compartmental interaction. To test this, we run simulations without loop extrusion, but instead we lowered the B-B attraction until the degree of compartmentalization was reduced as much as by adding loop extrusion. We find, however, that Hi-C maps (Fig. S7) and the compartment profile autocorrelation (Fig. 5A) behave differently. Indeed, for reduced B-B attraction we see little evidence of compartment coarsening, i.e. loss of shorter compartment regions, as the autocorrelation didn’t change considerably. We thus conclude that the impact loop extrusion on compartmentalization can not be described by a reduction compartmental interaction.

We thus asked if the non-equilibrium nature of loop extrusion is essential for its interference with compartmentalization. Indeed, loop extrusion is a non-equilibrium active process because loops start small, grow, and then are released when the loop extruder dissociates. Two aspects of this process can interfere with compartmentalization: (i) extruded loops connect loci irrespective of their compartmental identity, and (ii) the active nature of loop extrusion can increase mixing of compartments because loci need some time to re-segregate after being brought into contact by active loop extrusion. To examine the relative contributions of these factors we compare dynamically *growing loops* and *static loops*. We choose an ensemble of static loops from simulations with loop extrusion, but now loops remain static while the chromatin fiber is subject to thermal motion (see Supplement for details). We find that TADs are still visible in the corresponding Hi-C maps, albeit weaker (Fig. 5B), but that the degree of compartmentalization is almost as strong as when loops are completely absent (Fig. 5C). Also the compartment profile autocorrelation for static loops resembles the one without loops (Fig. 5A). In order to generalize the dichotomy of static *vs* dynamic loops we varied the loop extrusion speed and found that compartmentalization decreases for faster LEFs (Fig. 5C, and Fig. S6). Taken together, our results suggest that static loops contribute little to the observed compartment mixing, indicating that the non-equilibrium nature of active loop extrusion is central to interference with compartmentalization.

The non-equilibrium effect of active loop extrusion can be further strengthened by entrapment of the fiber in the dense network of chromatin surrounding it (52, 53). It is generally known that the amount of chain passing, which is enabled by topoisomerase II activity in the cell nucleus, has a great influence on relaxation times of polymer systems (37, 38). We thus alter the stringency of such topological constraints by changing the repulsive core of the monomer interaction potential E_rep_. We find that more stringent topological constraints reduce compartmentalization (Fig. S8), and that the loop extrusion impact on compartmentalization increases (Fig. 5D). Thus, our findings suggest that loop extrusion keeps chromatin far from equilibrium, with topological constraints reinforcing this effect.

The non-equilibrium nature of loop extrusion not only leads to compartmental mixing, but also directly affects non-compartment related quantities that can potentially be addressed experimentally. In particular, we consider the 3D size of an extruded loop, as measured by its radius of gyration *R*_*g*_ (Fig. 5E). We find that actively extruded loops are more compact than static loops, and that the compaction increases with loop extrusion factor speed (Fig. 5F, see Supplement for details). This is expected, because loci that are brought into proximity by loop extrusion need time to move apart by thermal diffusion (Rouse diffusion, Fig. 5E). Finally, we ask how active loop extrusion is reflected in the overall dynamics of the chromatin fiber by measuring its mean square displacement (MSD). Specifically, we asked if loop extrusion could be understood as an increased effective temperature, a conceivable consequence of the energy input from molecular motors. We find, however, that the MSD is elevated only on the time scale of loop extrusion without affecting the displacement on longer times (Fig. 5G). This is inconsistent with an elevated effective temperature, which would increase MSDs uniformly. In conclusion, we found that neither (i) elevated effective temperature, nor (ii) static or very slow loops, nor (iii) reduced compartmental interaction can reproduce the effects of loop extrusion, which underlines that it is a true non-equilibrium effect that can be thought of as active mixing of the polymer system. Experimental ramifications of these findings are discussed below.

**Fig. 5.**
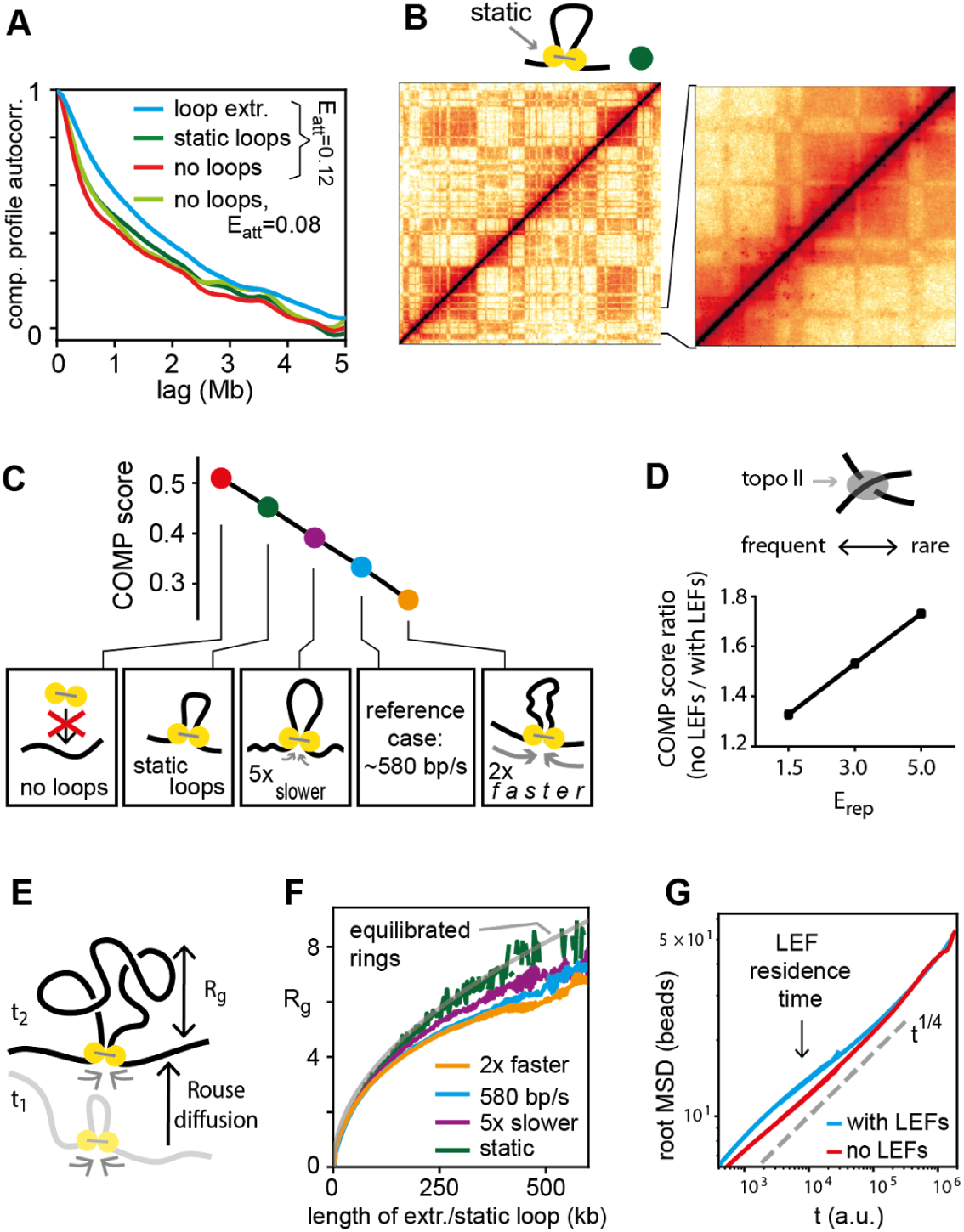
The non-equilibrium nature of loop extrusion. (A) Compartment profile autocorrelation as in Fig. 2A, compared with static loops and reduced B-B attraction. (B) Simulated Hi-C map for a chromatin fiber decorated by static, instead of extruding loops. Static loops have the same statistical properties as extruding loops studied above. (C) Degree of compartmentalization as a function of LEF speed. (D) The impact of loop extrusion on compartmentalization, measured by the ratio without/with loop extrusion, increases for reduced chain passing, i.e. reduced topoisomerase II activity (parametrized by E_rep_, the repulsive part of the monomer interaction potential). (E) Length scales relevant for equilibration of a loop: radius of gyration *R*_*g*_ and diffusional displacement during loop growth. (F) *R*_*g*_ follows the equilibrium theory (grey) for static loops, while with increasing LEF speed loops are more compact. (G) MSD of chromatin with/without loop extrusion differs on the LEF residence timescale, but not globally, indicating that loop extrusion can not be described as an elevated effective temperature.

### Changes in TADs and compartmentalization can reveal the mechanism underlying those changes

To consolidate our results we consider how the strengths of TADs and compartments are connected to each other, and how they can be altered by biological perturbations at the molecular level. To this end we measure how the strengths of TADs compartments change as we vary (i) the characteristics of the loop extrusion machinery, (ii) impermeability of boundaries to extrusion, (iii) the strength of epigenetically encoded compartmental interaction, and (iv) nuclear volume (Fig. S10). We find (Fig. 6) that alterations of the loop extrusion process, namely of the residence time of LEFs, their linear density and the speed of extrusion, resulted in coordinated and opposite changes in TADs and compartmentalization: higher activity of the loop extrusion leads to stronger TADs and weaker (more mixed) compartments, and vice versa. Interestingly, simulated activation or inhibition of topoisomerase II, allowing more or fewer chain passings, show a similar trend. Alteration of the boundary strength, however, shows a different pattern; it strongly affects TADs but leaves compartmentalization almost unaffected (as loop extrusion is preserved, see CTCF removal above). Strikingly, when nuclear volume or the compartmental interaction (i.e. B-B attraction) is changed we observe a third type of behavior: changes in compartmentalization but not in the strength of TADs.

This analysis provides a new approach to interpreting existing and future experimental data, suggesting that coordinated changes in TADs and compartments reflect changes in the loop extruding machinery of cohesin or topoisomerase II activity, changes in TADs that leave compartments unaffected most likely come from altered insulation (boundary proteins such as CTCF, and potentially YY1 and Znf143, either globally or from perturbing particular sites); and changes in compartments that do not affect TADs reflect changes in nuclear volume or the epigenetic landscape of histone modifications or the molecules that mediate their interactions.

**Fig. 6.**
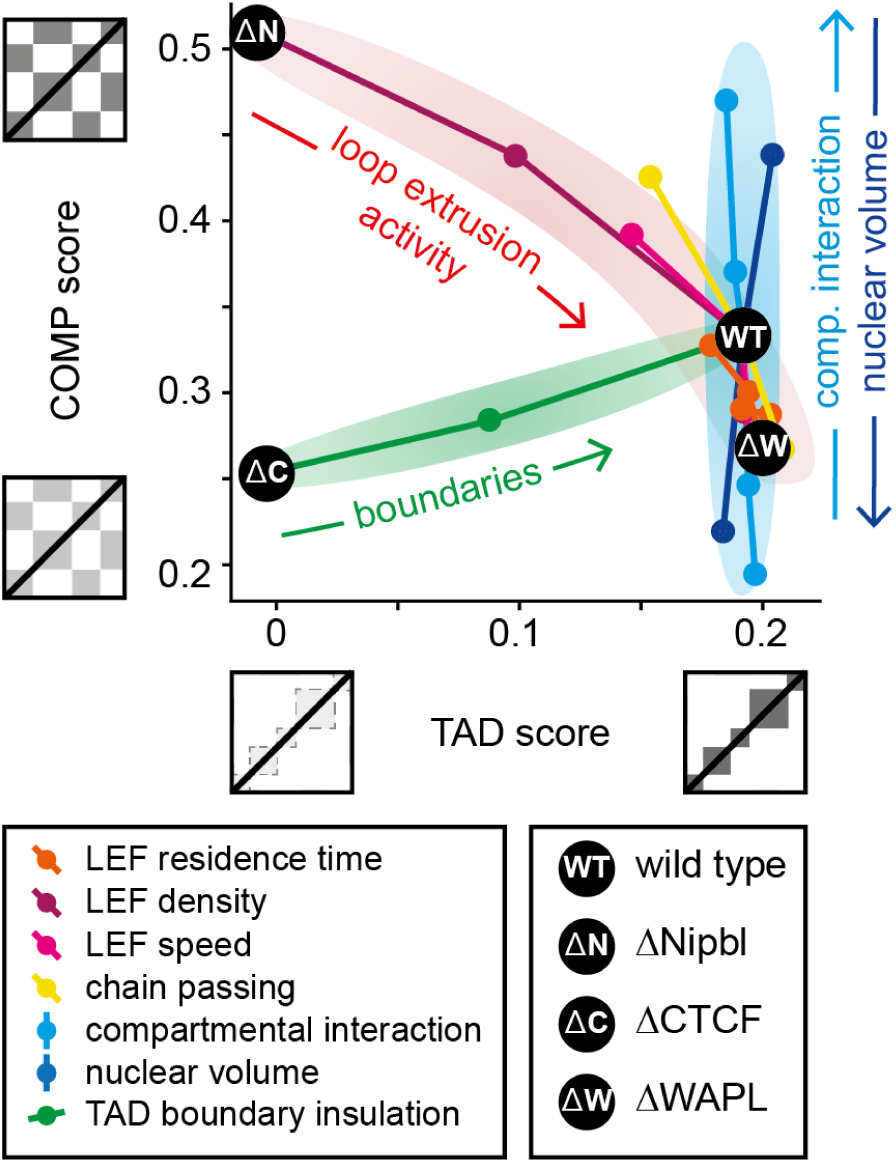
The joint variation of TAD and compartment strength indicates the underlying parameter change. Three main classes of parameters are identified: a tradeoff between compartmentalization and TADs is observed for parameters related to cohesin dynamics and for the frequency of chain passing (topoisomerase II activity), compartmental interaction and nuclear volume mainly affects compartments, and the permeability of TAD boundaries mainly affects TADs. Complete lists of parameters for all data points are given in the Supplement (Fig. S9). The black dots indicate our simulations of wild type cells as well as removal of cohesin (by Nipbl deletion), of CTCF and of Wapl.

## Discussion

We have elucidated a key step towards a complete model of interphase chromatin: the interplay of loop extrusion and compartmental segregation, two mechanisms that shape major features of chromosome organization in mammals. Motivated by recent experiments that point toward such an interplay (11, 14, 36), we used polymer models of chromosomes to investigate whether simultaneous action of loop extrusion and compartmental segregation can quantitatively reproduce experimental findings. We found that this is indeed the case for all three perturbations, namely removal of chromatin associated cohesin by Nipbl removal, removal of the TAD boundary protein CTCF, and removal of the cohesin unloader Wapl. The key insight is that loop extrusion counteracts compartmental segregation. This argues against a hierarchical organization which claims that TADs are building blocks of compartments, and replaces it with a more complex picture where the active loop extrusion partially overrides innate compartmentalization preferences.

Specifically, we found that (i) removal of the cohesin loader Nipbl reveals the intrinsic compartment structure because segregation is no longer suppressed by loop extrusion. (ii) Removal of the loop extrusion boundary element CTCF removes TADs because loop extrusion is not confined to specific domains, but continues to locally compact chromatin and to counteract compartmental segregation. (iii) Removal of the cohesin unloading factor Wapl increases cohesin residence time on DNA and thereby leads to longer and more abundant loops, which at the same time strengthens TADs and weakens compartmentalization due to enhanced compartment mixing.

Our mechanistic model relies on simplifying assumptions that we now address. First, the microscopic biophysical mechanisms that drive compartmental segregation remain unknown. Here, we assumed a phase separation process, in line with experimental indications for heterochromatin formation (44, 45), and we furthermore postulated a specific short range attraction between chromatin loci of type B. It is important to emphasize that this constitutes a minimal model for compartmental segregation. Other interaction potentials or even different mechanisms of segregation could be present as well. For example, segregation based on differences in activity instead of contact interaction is a plausible scenario (54–56). Similarly, one may want to model the role of interactions between heterochromatin (B type regions) and nuclear lamina. Our focus on B-B interactions is motivated by the observations that cells lacking naturally or artificially lamin and/or B receptor, such as rod cells nevertheless exhibit compartmentalization (Falk et al., in preparation).

Similarly, the microscopic mechanisms behind loop extrusion remain highly enigmatic and controversial. In particular, processive motion has been demonstrated in *in vitro* only for condensin (28) while corresponding evidence is still missing for cohesins, which are relevant in interphase. Furthermore, experiments are at odds with a simple picture where the sole function of Scc2/Scc4 complex is to facilitate cohesin loading while Wapl determines its residence time on chromatin, and rather suggest that Scc4 also regulates the processivity and/or the residence time of cohesin on DNA (14), that Wapl/Pds5 assists in loading and unloading (57), and that transcription plays a major role in positioning cohesins (58). Consequently, several parameters in our mechanistic model of loop extrusion are known with limited accuracy. Those include the number of DNA bound loop extruding factors, their processivity, their speed, details about the extrusion process (e.g. one-sided vs two-sided), and interaction with other proteins like CTCF, Nipbl, Wapl and Pds5 (13). In light of such uncertainties we use simulations to establish *consistency* of our mechanistic model with experimental observations.

Surprisingly, our relatively simple and general mechanistic model was able to achieve consistency with experiments reproducing a number of features, such as TADs, compartmentalization, and the contact probability *P(s)* curves, for a diverse set of unrelated experimental perturbations. In the future, an iterative process of increasingly specific experiments and more constrained simulations will show how far the loop extrusion and compartment segregation model can go in quantitatively explaining chromatin organization.

We finally discuss experimental ramifications and potential tests of our model. While our study was motivated by specific alterations of the loop extrusion machinery (namely abundance, boundaries, and processivity), our results go beyond explaining these experiments and make specific predictions. In particular, experiments where the speed of loop extrusion factors or topoisomerase II activity is altered are expected to see a tradeoff between TAD strength and compartmentalization. Conversely, perturbations altering the nuclear volume or the compartmental interaction, e.g. by changing the epigenetic landscape or mediators of compartment interactions, possibly HP1 (44, 45), are expected to affect compartmentalization, while leaving TADs unaffected. Furthermore we showed that when faced with an experimental phenotype for which the underlying microscopic alteration is not known, the joint variation of TADs and compartmentalization can help to unravel it: variations in TAD strength alone indicate that only TAD boundaries are affected, variations in compartmentalization alone indicate that the compartmental interaction is changed, while a tradeoff between TAD strength and compartmentalization stems from changed cohesin dynamics or topoisomerase II activity. As an example, a recent comparison of maternal and paternal pronuclei demonstrated similar TAD strength, but considerably weaker compartmentalization in maternal zygote; our results here suggest that this is due to differences in the epigenetic landscape, and possibly a lack of heterochromatin of those pronuclei (10). Finally, we found that characteristics of the 3D path of chromatin bear information about specific aspects of loop extrusion: loops are more compact in 3D space when extrusion is fast, consistent with the observation that changing extrusion speed can disentangle contact frequency from average spatial distances (59). As high resolution imaging of chromatin is making dramatic progress (7, 60, 61), such questions may be addressed in the near future.

In conclusion, our work shows that the interplay of active loop extrusion and compartmental segregation shapes chromosome organization in interphase. More broadly, we hope that the principle that active processes can oppose equilibrium energetics, can serve as a paradigm for future biophysical research.

## Methods

Our study relies on coarse grained molecular dynamics simulations of chromatin subject to loop extrusion and compartment segregation. Simulations were performed based on OpenMM (62, 63). In brief, our approach is to generate a large number of polymer conformations from which a simulated Hi-C experiment produces contact maps that are compared with experimental Hi-C data. We typically simulated a 20,000 monomer chain, with one monomer corresponding to 2.5 kb. The TAD structure was defined by random positioning of boundary elements along the polymer. The average TAD size was 375 kb (150 monomers). Compartments were also placed randomly and not correlated with TADs. We used a randomly generated TAD and compartment structure because there is no uniquely agreed upon method for calling them from experimental data and because our results on aggregated quantities like degree of compartmentalization, compartment profile autocorrelations and contact probability scaling can be equally well made with random TADs and compartments. Loop extrusion factors are implemented as a bonds between not necessarily adjacent monomers. When an LEF takes a step from, say, monomers (i,j) to (monomers (i-1,j+1) the old bond is deleted and is replaced with a new bond. Details are given in the Supplement.

## Acknowledgements

We gratefully acknowledge funding from the following grants: NSF 1504942 (Physics of Chromosomes) and NIH GM114190 (Polymer Models of Mitotic and Interphase Chromosomes) to LM; and support of the 4D Nucleome NIH Initiative DK107980 (Center for 3D Structure and Physics of the Genome).

## Supplemental Information

### Polymer simulations

Polymer simulations were performed using a lab written wrapper (available at https://bitbucket.org/mirnylab/openmm-polymer) around the open source GPU-assisted molecular dynamics package Openmm (62, 63). Polymers are represented as a chain of monomers with harmonic bonds, a repulsive excluded volume potential, and an additional small attraction (for the interaction of two monomers of type B, see main text). Loop extrusion is modeled as an additional harmonic bond between two not necessarily adjacent monomers (i,j). When the loop extrusion factor takes a step this bond is deleted and replaced by a bond between (i-1, j+1), provided that there are no loop extrusion boundaries at the updated sites. Details of the loop extrusion simulations can be found in (16).

For each Hi-C contact map simulations were carried out as follows. Unless otherwise mentioned, we use a chain 20,000 monomers (equivalent to 50 Mb) in periodic boundary conditions and a volume density of 0.2 monomers per unit volume. Simulations were initiated with compact, unknotted chain conformations and allowed to expand before conformations were recorded. Expansion times were chosen such that the mean square displacement of a monomer equals the radius of gyration of the polymer coil (for periodic boundary conditions) or the confinement volume radius (for spherical confinement). Subsequently, the simulation was run for the expansion time. During this part 500 conformations were recorded. The entire procedure was repeated 20 times, yielding 10,000 conformations. From these, a Hi-C map was computed by defining a cutoff radius, mimicking the crosslinking radius in an actual Hi-C experiment. The cutoff radius was four monomer diameters (we verified that result are insensitive to the cutoff).

### Converting simulation parameters to real time and length scales

In our simulations, the chromatin fiber is a chain of monomers connected by harmonic potentials subject to a monomer interaction potential (excluded volume and B-B attraction) and bending stiffness. Furthermore, a 1D layout of LEF boundaries and compartments is defined along the polymer, which involves sizes of TADs and compartments. These simulation parameters need to be converted to parameters of the actual chromatin fiber in the cell. We here address this conversion, noting up front that many involved quantities are not well characterized. Our conversion should therefore be taken only as a rough guide.

We start by considering length scales. The spatial coarse graining of our simulations, i.e. how many base pairs correspond to one monomer in our simulation, is chosen by computational feasibility considerations. Conversion to physical parameters proceeds in the following steps: 1.) choice of the persistence length for the simulated chromatin fiber, 2.) estimation of the persistence length of the biological chromatin fiber, and 3.) determining conversion factors from comparing 1.) and 2.).

**1. persistence length in simulations**

We choose *l*_*p*_ = 2 monomers.

**2. persistence length of the biological chromatin fiber**

The persistence length of the fiber can be given either in nm or in bp, related by the compaction ratio c, which is measured in bp/nm. Reported values, based on cyclization experiments (1, 64) or imaging (65), vary greatly and are roughly in the ranges *c* = 25…150 bp/nm and *l*_*p*_ = 30…200 nm. For our conversion we use *c* = 50 bp/nm and *l*_*p*_ = 100 nm. During preparation of this manuscript we learned of an estimate based on parameter sweeps for whole nucleus simulations in yeast (66) arriving at *c* = 61 bp/nm and *l*_*p*_ = 88 nm.

**3. Comparison of simulation and biological parameters**

With the above parameters for the persistence length and compaction ratio we arrive at the following conversion

1 monomer = 50 nm

= 2.5 kb

We point out again that this is a rough estimate only, due to the experimental uncertainty of the compaction ratio and the chromatin persistence length. Our values are very close to the estimate from (67) which gave 53 nm for 3 kb.

Next we aim at converting time scales. First we note that, even with spatial coarse graining, accurate molecular dynamics simulation of our system is computationally not feasible for times we are interested in (at least several minutes). This is true even in implicit solvent simulations. Our approach is to strongly reduce the collision frequency with the implicit solvent. This comes at the price of unrealistic short-time dynamics: the ballistic flight length of our monomers becomes unrealistically large, namely it is of the order of the monomer size. Yet, on times considerably longer than the collision frequency, Rouse diffusion is recovered for our polymer. We thus need to convert simulation time units to physical units. This is based on **comparing MSDs**, namely the relation between simulation and experiment, where *D* is the anomalous diffusion constant. As the length conversion is known from above, this fixes time scale conversion.

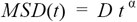

In simulations, *D* depends on several simulation parameters, in particular on the density of the implicit solvent (parametrized by the “thermostat” value in openMM) and is obtained from simulations without loop extrusion.

For the biological case, values for *D* have been reported as *D* = 0.01 μ^2^/*s*^1/2^ in yeast (68) and *D* = 0.015 μ^2^/*s*^1/2^ in mammals (69). Note, however, that these already inculude active processes, including loop extrusion. We thus also consider a smaller value of *D* = 0.005 μ^2^/*s*^1/2^. As a reference case we here chose *D* = 0.01 μ^2^/*s*^1/2^.

For a chosen pair of *D_exp* and *D_sim* we determine the conversion factor according to the formula

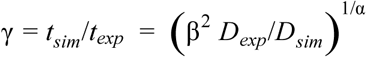

where β is the length scale conversion from above and α is the time exponent, for which we use 1/4 in accordance with experiments and simulations.

### Choice of simulation parameters for TADs, compartments and loop extrusion factors

#### TADs

TADs are defined by their boundary positions (CTCF sites) and the boundary insulation strength. As we do not aim to reproduce a particular genomic region we place TAD borders randomly along our polymer, taking only the average genomic distance between TAD boundaries from experimental considerations. To this end we analyzed annotated domains and loops from (27). Loops had an average size of 1.1 Mb. This, however, included a few extremely large loops. Loops shorter than 5Mb had an average size of 387 kb. Domains had an average size of 258 kb. For convenience, we chose an average TAD size of 375 kb (150 monomers) in our simulations. We draw TAD sizes from a distribution that has an exponential tail but also suppresses very short TADs, namely *p*(*d*)= *d*/*d*_0_ *exp*(- *d*/*d*_0_) which has a mean of 2*d*_0_ which we chose to be 375 kb. The mean TAD size of the actually chosen TADs came out as 402 kb (161 monomers).

TAD boundaries are characterized by their insulation strength (or, conversely, permeability) to LEFs. As mentioned in the main text, the exact values are not known. We chose an insulation of 90 %. We point out, however, that the exact value is of minor importance (see Fig. S5 for a parameter sweep).

#### Compartments

Our goal was to create compartments that are typically larger than TADs but are interspersed with small compartments of the respective other type. To that end we proceed as follows: We draw compartment sizes from a distribution with mean c1. After each draw, with probability *p* we interst a small interstition by drawing a compartment size from a distribution with mean c2. This has the effect that, when a small interstition is inserted, the next larger compartment is of the same type as the previous larger compartment. We chose c1=100, c2=10 and p=0.7 for our chains with 20,000 monomers and p=0.6 for chains with 10,000 monomers (Fig. S2).

The choice of these parameters were guided by considering the compartment profile autocorrelation. In particular, we wanted to achieve a decay length that (i) resembles the decay length of compartment profiles from experimental Hi-C maps, and (ii) exhibits a change upon removal of loop extrusion that is similar to experiments (see also section “Discussion of quantitative differences between simulations and experiments for Nipbl removal” below).

#### Loop extrusion factors

Loop extrusion factors (LEFs) are characterized by their density on the fiber, their processivity, and their speed. As mentioned in the main text, we chose a processivity λ =250 kb and an average separation of *d*=750 kb. This is a factor of 2-6 lower than suggested by a previous study (16). We chose these parameters, however, because they lead to better consistency with the experimental perturbations studied here. In particular, we took into consideration: (i) the compartment profile autocorrelation change upon removal of loop extrusion, (ii) the very small change in contact probability scaling upon removal of CTCF, and (iii) substantial reduction in compartmentalization in removal of WAPL. As a full simulation of a relatively large system (20000 monomers, 10000 conformations) is required for each set of parameters for each layout of TADs and compartments we refrained from an extensive sweep of all possible parameters and rather were guided by a hand-picked sampling of parameter space. Furthermore, We stress that our major focus here is not to tweak simulation parameters to match experiments as much as possible, but rather to demonstrate that the interplay of loop extrusion and compartmental segregation predicts several observables correctly across different experimental perturbations.

As active loop extrusion is a non-equilibrium process, the LEF speed *v* matters: for slow enough LEFs the loops are equilibrated by thermal diffusion while fast LEFs lead to non-equilibrated loops. It is currently not possible to determine which regime is realistic, because *in vivo* LEF speeds are unknown and thermal diffusion of the chromatin fiber in the absence of LEFs is insufficiently characterized. Rough estimates are: (i) TADs of up to 1MB in size are observed, while LEF residence times are of the order of 1hr, suggesting a speed of at least ~278 bp/s. (ii) Data for bacterial condensin suggests 770 bp/s (29). (iii) *In vitro* experiments for condensin suggest 63 bp/s (28). We here chose a reference case that corresponds to approximately 577 bp/s. As this value is subject to considerable uncertainty we study the dependence of our results on the LEF speed in the main text.

We here give a conversion of our simulation LEF speeds to actual speeds, based on the conversion of length and time scales discussed above.

**Table S1.** Conversion of LEF speeds in simulation to physical units. Note that this conversion is based on several parameters that are known with insufficient accuracy as discussed above (section “Converting simulation parameters to real time and length scales”). Most notably, as shown in this table, the conversion depends sensitively on the (anomalous) diffusion constant for the chromatin fiber in the absence of loop extrusion. The values in bold face are what we consider as reference case in the main text.

| assumption on $D_{exp}[\mu^2/s^{1/2}]$ (chromatin) | simulation LEF speed (3D steps / 1D step) | physical LEF speed (bp/s) | comment |
| --- | --- | --- | --- |
| 0.005 | 100 | 288 | 2x faster |
|  | 200 | 144 | reference case |
|  | 1000 | 29 | 5x slower |
| **0.01** | 100 | 1154 | 2x faster |
|  | **200** | **577** | **reference case** |
|  | 1000 | 115 | 5x slower |
| 0.015 | 100 | 2596 | 2x faster |
|  | 200 | 1298 | reference case |
|  | 1000 | 260 | 5x slower |

### Definition of compartment and TAD scores

A compartment score quantifies compartmentalization of a Hi-C contact matrix. Such a score should measure the excess of contact between monomers within a compartment over across a compartment. This required a definition of which monomers belong to which compartment. Two different approaches can be taken: Either monomers are classified based on their underlying properties, or based on their (average) environment in 3D space. These classifications need not be equivalent. Indeed, we have shown in this paper that in particular loop extrusion can counteract segregation of small segments into their native compartments.

In simulations we know for each monomer if it is of type A or B and we can thus use this information to develop an appropriate compartmentalization score (COMPscore1 below). The same is not true for experimental data, where the underlying types of loci are not known. We thus use a different method for experimental data, consistent with previous convention (11, 36), namely COMPscore2.

### COMPscore1: Inferring compartmentalization from simulated data

This score assumes that we know the compartmental type of each monomer. We can use this score for Hi-C maps from simulations, where the types are known by definition. COMPscore1 is computed as follows: for a given distance *s* along the polymer we consider all pairs *(i,j)* with *j-i*=*s*, i.e. we consider a diagonal with distance *s* from the main diagonal. For a given *s* we count the the number of pairs *(i,j)* where *i* and *j* belong to the same compartment, *#pairs_within_comp(s)*, and to two different compartments, *#pairs_across_comp(s)*. Furthermore, we count the actual number of contacts in the Hi-C map *hmap* such that *i* and *j* belong to the same compartment,

*#contacts_within_comp(s) = sum(hmap(i,j)* if *i* and *j* from same compartment and *j-i*=*s)* and analogously for different compartments. From this we define

*av_within(s)* = *#contacts_within_comp(s)* / #pairs_within_comp(s), which is the average value of the diagonal with distance *s* from the main diagonal of *hamp* restricted to pairs that belong to the same compartment. In the same way we define *av_across(s)* where the two loci belong to different compartments. From this we compute

*COMPscore1(s)* = (*av_within(s) - av_across(s)) /* (*av_within(s) + av_across(s)).*

This yields a value between −1 and 1, where *COMPscore1(s)=0* if a contact between *i* and *j* is equally likely whether they belong to the same or different compartments, and *COMPscore1(s)=1* if there are only contact within compartments but none across, and *COMPscore1(s)=−1* if there are only contacts across compartments. Finally, we average this this measure over all *s* from 0 to L/2, where L is the size of the contact map:

*COMPscore1 = <COMPscore1(s)> for s = 0…L/2.*

The properties of this score include: (i) COMPscore1 is insensitive to the absolute number of reads in the matrix (the “sequencing depth”). More precisely, multiplying each entry of *hmap* by a factor does not change the score. (ii) COMPscore1 weighs all distances form the main diagonal equally (i.e. the “scaling” is unimportant); it measures the “contrast” along each diagonal, irrespective of the average value. (iii) COMPscore1 is between −1 and 1, with 0 for no compartment segregation and 1 for perfect compartment segregation.

### TAD score: Inferring TAD strength from simulated data

The TAD score is defined in complete analogy to COMPscore1. The only two differences are: (i) *#pairs_within_TADs(s)* and *#contacts_within_TADs(s)* are are counted within each TAD individually, there is no long-range association of TADs. (ii) The average across linear separations in *COMPscore1 = <COMPscore1(s)> for s = 0…L/2* is taken up to *L/2* where *L* is the size of the largest TAD in the system.

### COMPscore2: Inferring compartmentalization from experimental data

For COMPscore2, we first compute the “observed over expected” matrix from a given Hi-C contact matrix, i.e. each diagonal with distance *s* from the main diagonal is divided by the mean number of contact in this diagonal, which gives equal weight to all diagonals irrespective of their mean intensity. We then order rows and columns of the resulting matrix based on the compartment profile. This leads to accumulation of AA and BB contact in the quadrants along the diagonal and AB and BA contact in the off-diagonal quadrants (see Nora2017). We then compute

*COMPscore2 = (AA+BB) / (AB+BA),*

where AA is the number of contacts in the quadrant AA, etc.

COMPscore2 is not a value between −1 and 1, but rather always a positive number. Furthermore, *COMPscore2=1* if there is no compartmentalization and *COMPscore2>1* if there is. However, *COMPscore2* can be converted to a value between −1 and 1 in the following way

*COMPscore2_rescaled = (COMPscore2 - 1) / (COMPscore2 + 1).*

We apply this rescaling because ratios of *COMPscore2* are hard to interpret (e.g. the ratio of scores 1.01 and 1.1 is very close to unity, but the first has virtually no compartmentalization, the latter 10x more).

While we prefer to use *COMPscore1* for simulation data in principle, we use *COMPscore2_rescaled* for both experimental and simulated data when we compare compartmentalization between experiment and simulation.

Note that we don’t define a TAD score for experimental data as we refrain from quantifying TAD strength for experimental data in this study.

### Discussion of quantitative differences between simulations and experiments for Nipbl removal

We here address in detail quantitative discrepancies between our simulations and experimental data for the removal of cohesin/loop extrusion (Fig. dNIPBL).

(i) Our simulated compartments in the absence of loop extrusion appear somewhat smaller than in the experimental example, as seen in our HiC map as well as the more pronounced decay in compartment profile autocorrelation. We point out, though, that in many chromosomal regions the experimental compartments are also smaller (see Fig. S3 for an example), and that increased Hi-C resolution might reveal even finer structures. We also mention that the compartment profile autocorrelation is somewhat sensitive to the exact TAD and compartment configuration, not only to their average sizes. This is in line with differences between chromosomes as reported in (11).

(ii) Changes in contact probability scaling are more pronounced in simulations. It is noteworthy, though, that the characteristic loop extrusion hump is much more pronounced in many other datasets, including the ones used in this paper (see Figs. dCTCF, dWAPL).

(iii) The increase in compartmentalization upon removal of loop extrusion is somewhat more pronounced in simulations than in experiments. This could indicate that some compartment mixing remains present in experiments, either by residual cohesin (see Fig. S4 for 10% residual LEFs in simulation) or some other processes in the nucleus not considered here. We stress that our major focus here is to correctly qualitatively predict multiple observables across different experimental perturbations (see below for removal of CTCF and Wapl).

### Polymer simulations with static loops

In order to perform simulation with static loops our goal was to preserve the statistical properties of the loops from our reference case with active loop extrusion. We thus recorded loop configurations (left and right monomers (i,j) tethered by an LEF bond as described above) from our dynamic simulations at a given time point during the simulation. Those loops were put into our static loops simulation and the polymer dynamics were simulated as usual. In order to get better statistics of static loops we picked a new set of loop positions five times during the simulation and replaced the previous static loop bonds with the new ones. As our polymer equilibrates on the length scale of individual loops within the time between to successive recorded polymer configurations (out of the 500 recorded configurations for each run) this updating did not bring the polymer significantly out of equilibrium.

### Radius of gyration as a function of loop length

In Fig. 5F in the main text we plot the radius of gyration as a function of the length of an extruded or static loop. We here note that for an extruded loop its length increases over time. The length axes can thus be directly converted to a time axis, where time is measured since the attachment of the LEF. We also note that we only used loops that are not stalled (an LEF counts as stalled when at least one of the two sides stopped extruding because it bumped into a boundary element or another LEF). For short loop lengths (or times) the probability for an LEF to be stalled is very low. As loops get longer the chance to get stalled increases. As we are in the dilute regime, however, the chance to get stalled becomes substantial only when loops have grown to approximately their average separation between LEFs, i.e. 300 monomers (or 750 kb).

## Supplemental Figures

**Fig. S1.**
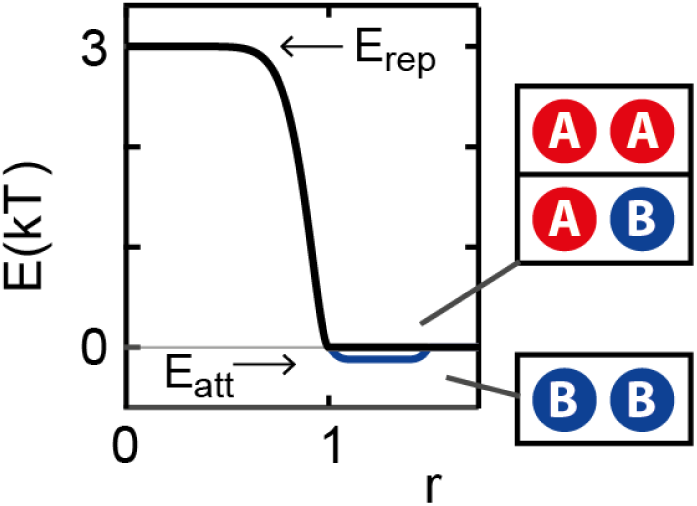
Monomer interaction potential for our polymer simulations of chromatin. Monomers of type B experience a slight additional attraction toward each other, which leads to spatial segregation of compartments. The depth of the attractive part is our parameter E_att_. It drives compartment segregation, and we refer to it as the compartmental interaction parameter.

**Fig. S2.**
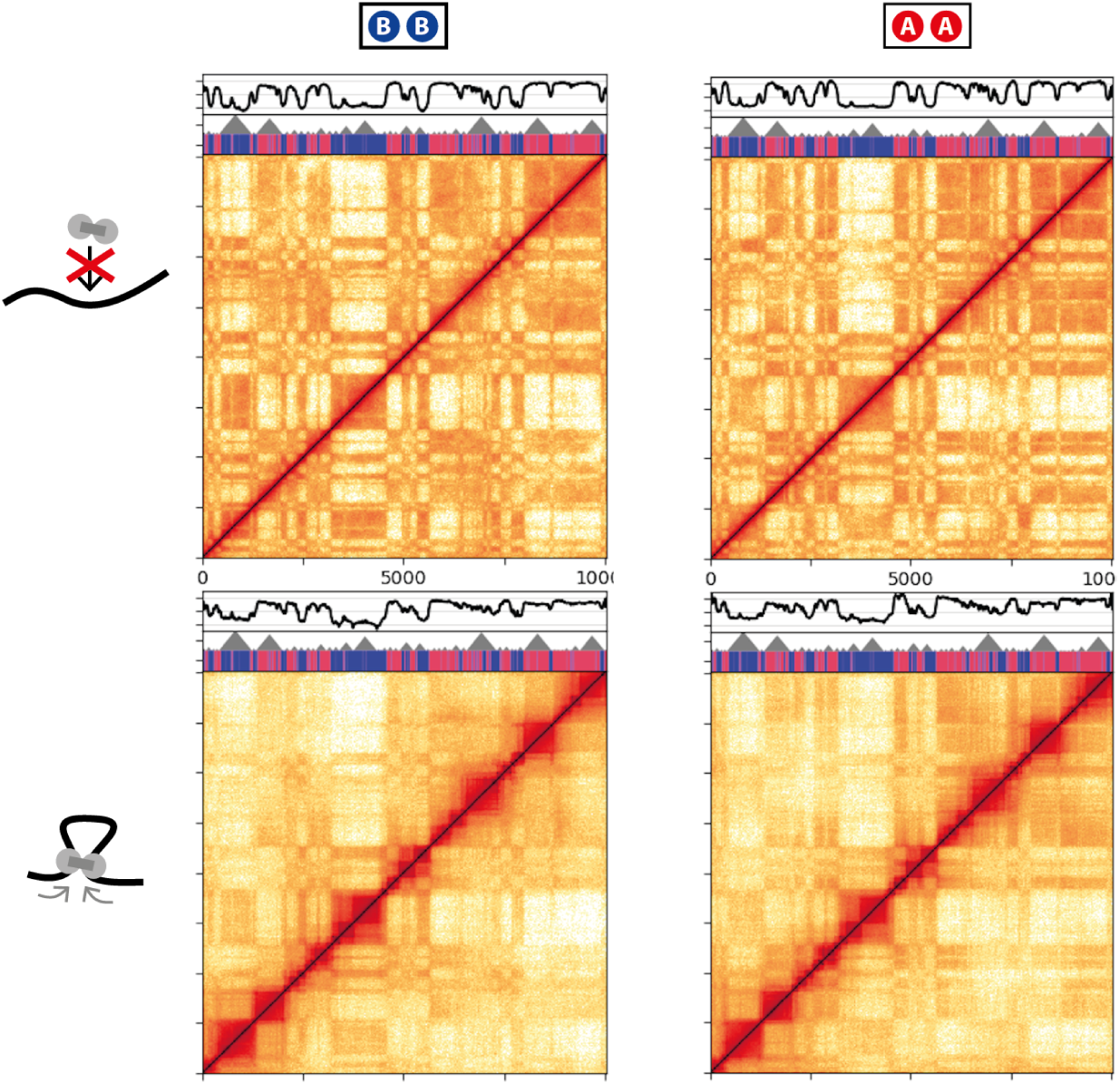
Contact matrices are insensitive to the details of the compartment segregation mechanism. In the left column segregation is induced by a slight attraction monomers of type B. This case was used throughout the main text. In the right column the attraction is between monomers of type A.

**Fig. S3.**
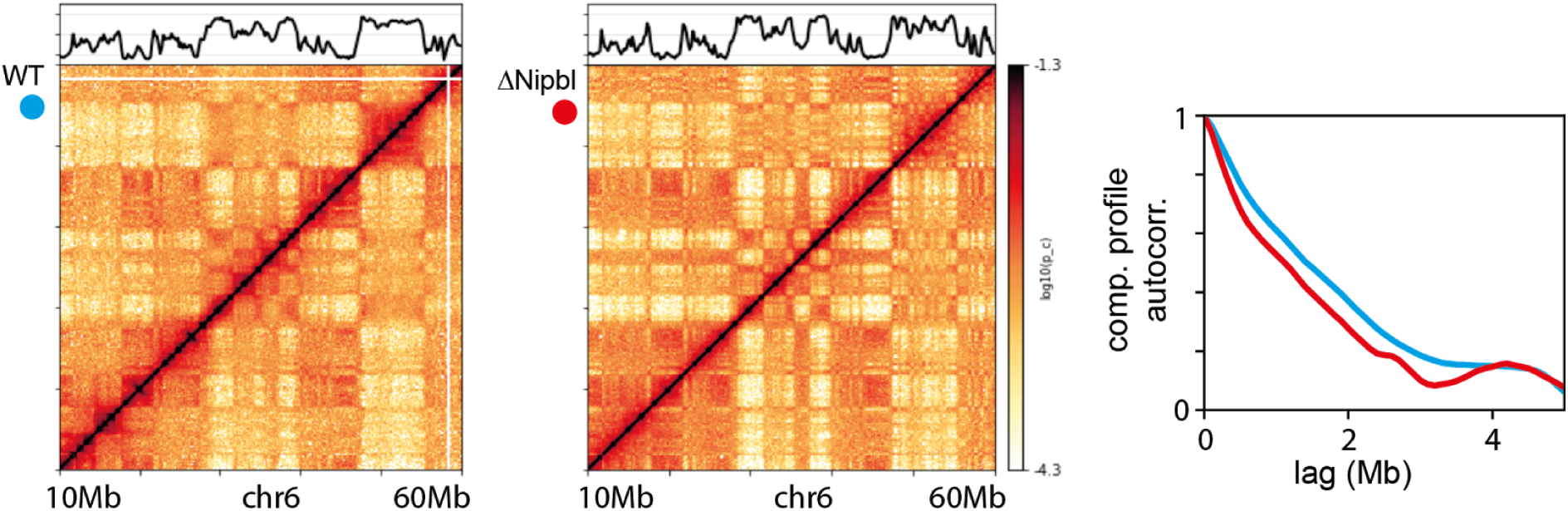
An example for a region with much smaller compartments than the ones shown in Fig. 2 in the main text. Data from (11).

**Fig. S4.**
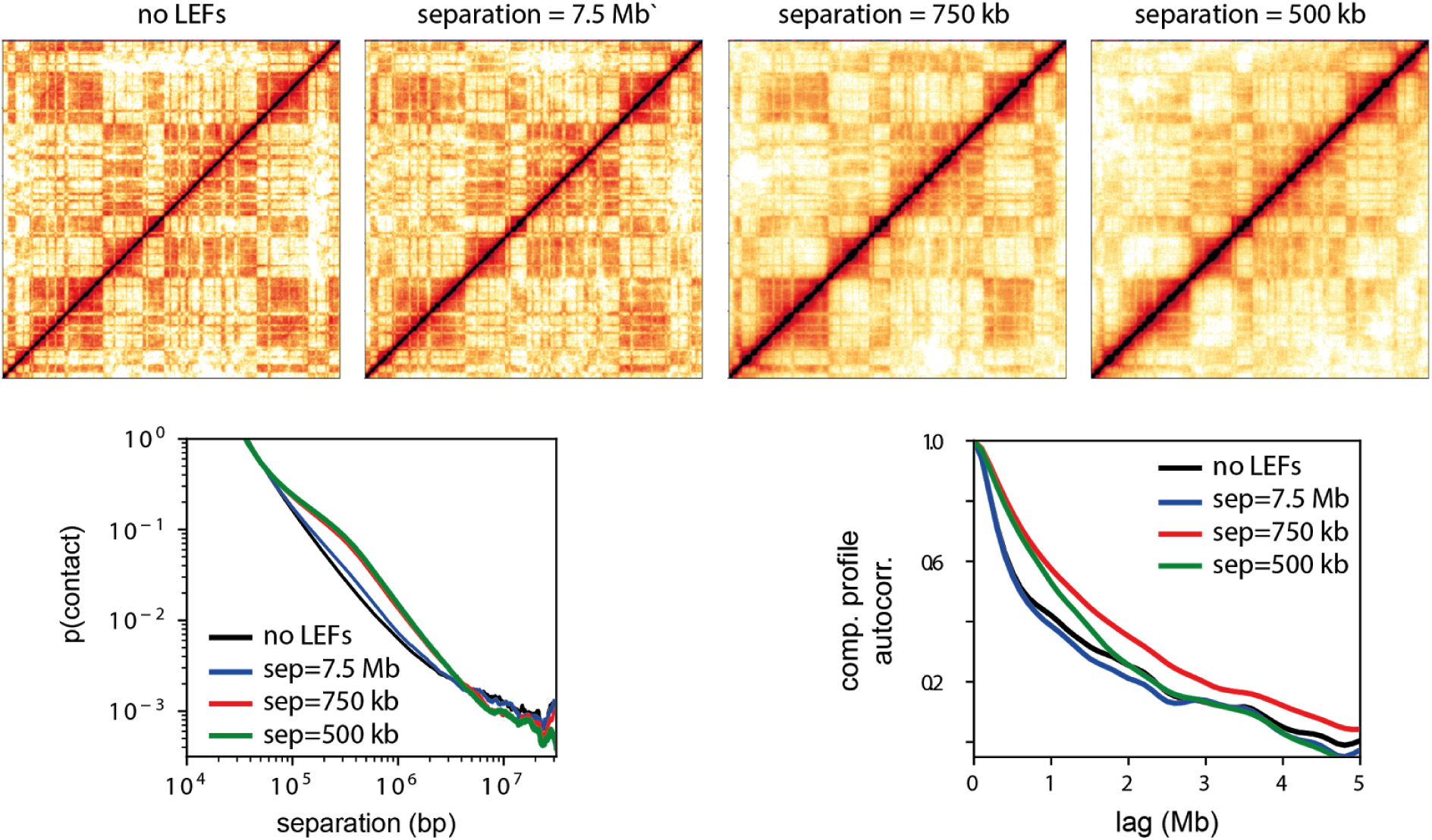
Contact matrices, contact probability scaling, and compartment profile autocorrelation for different LEF densities. All other parameters are as in our reference case in the main text.

**Fig. S5.**
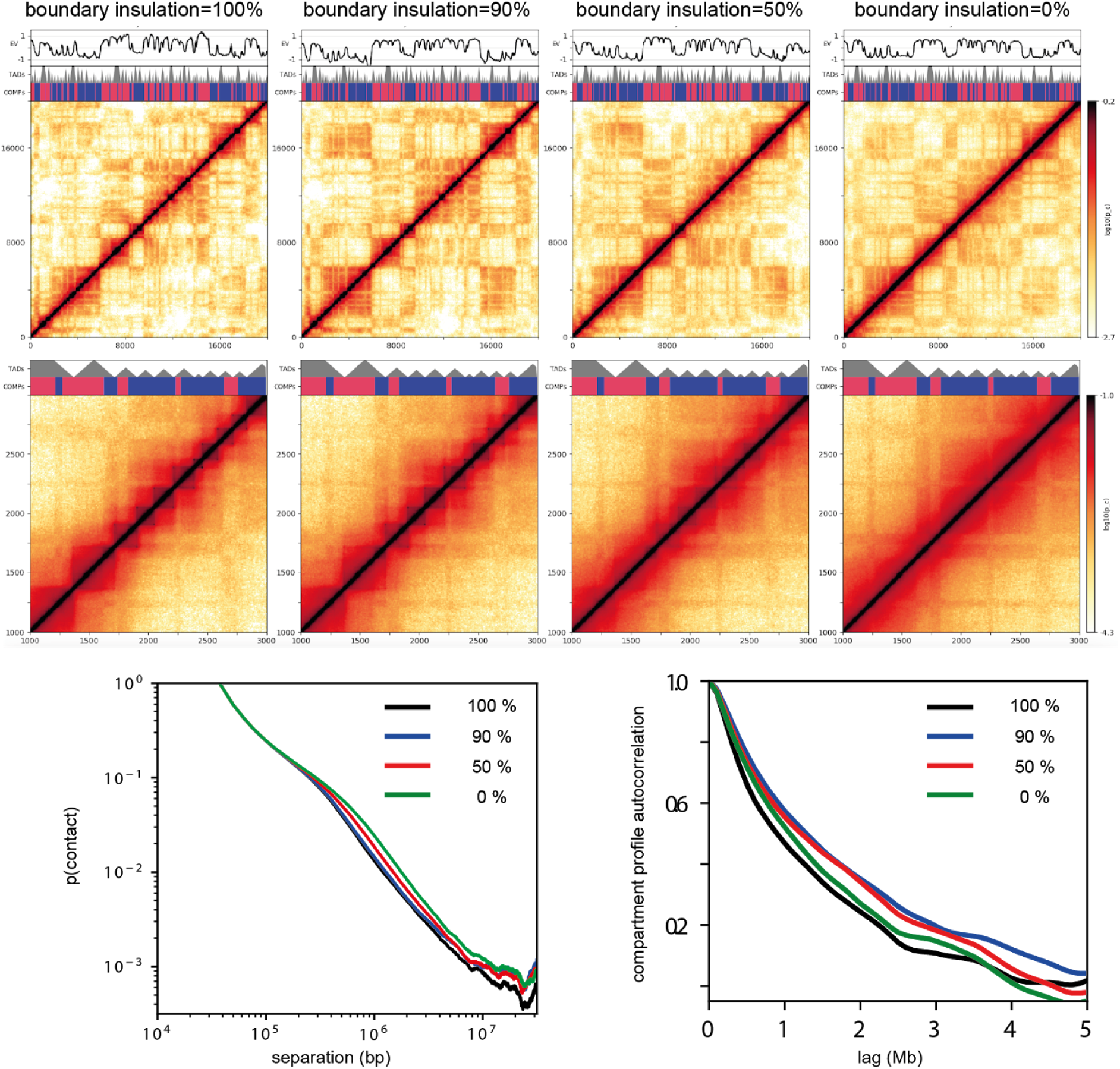
Contact maps for our 50 Mb chromatin fiber (top row) and a 5 Mb zoom in middle row), as well as contact probability scaling and compartment profile autocorrelation (bottom row) different boundary insulation strengths.

**Fig. S6.**
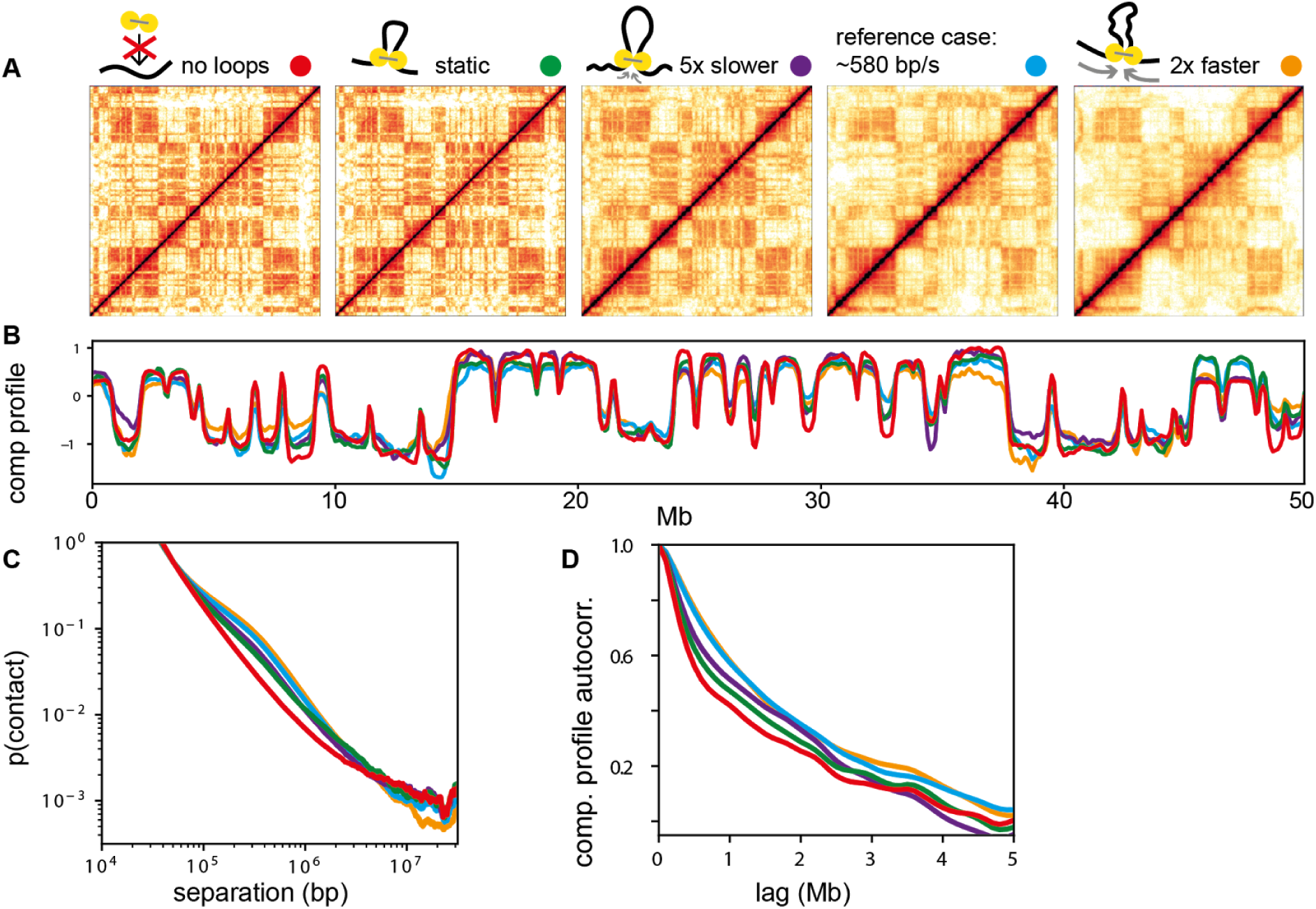
Effect of loop extrusion factor speed. Contact matrices (A), compartment profiles (B), autocorrelation thereof (C), and contact probability scaling (D) for varying loop extrusion factor speed, including static loops and absence of loops.

**Fig. S7.**
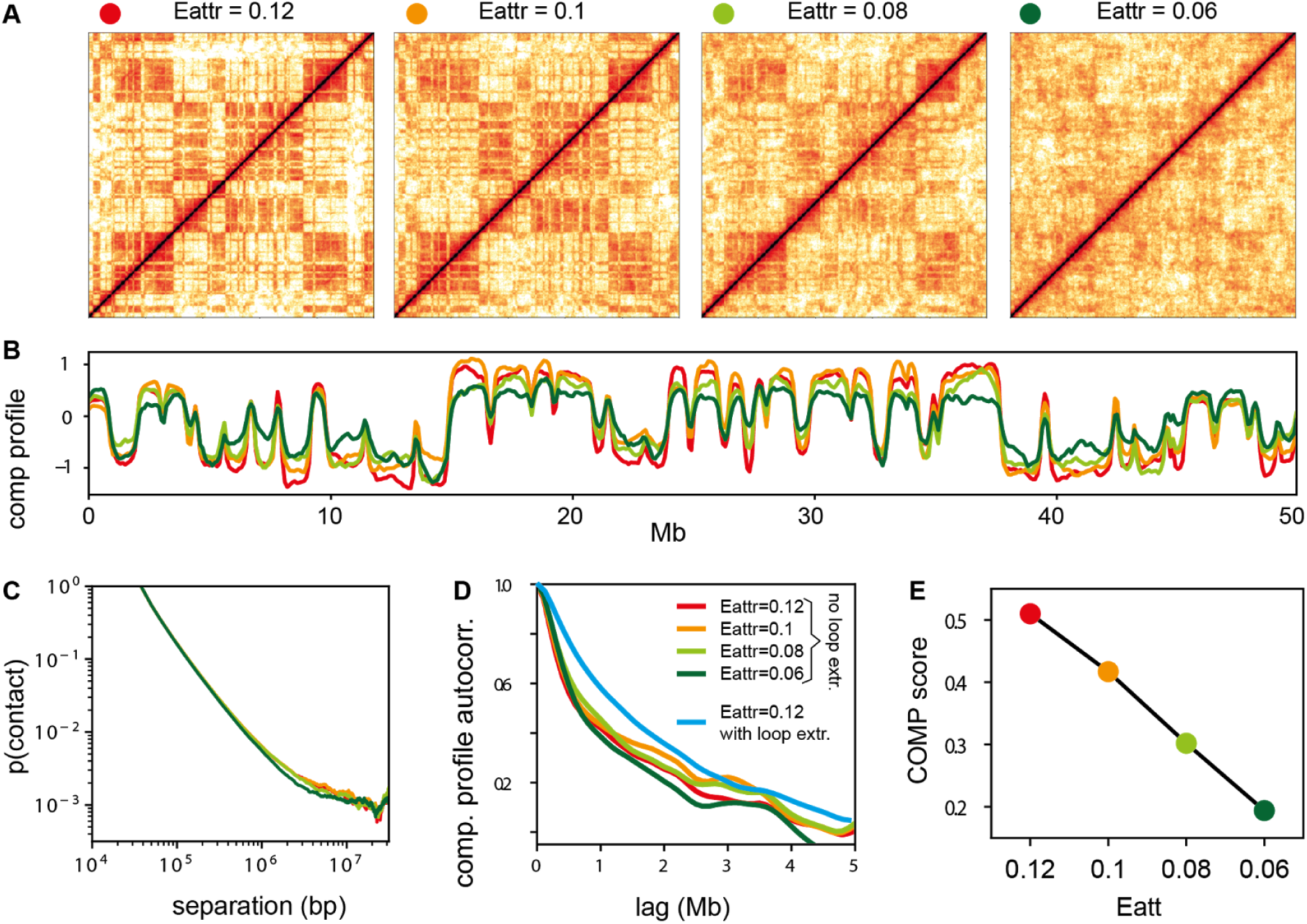
Effect of the compartmental interaction parameter E_att_. Contact matrices (A), compartment profiles (B), autocorrelation thereof (C), contact probability scaling (D) and degree of compartmentalization (E) for varying depth of the attractive part of B-B interaction potential (see Fig. S1 for its definition). In (D) the case with loop extrusion is also shown; it highlights that the effect of reduced compartment segregation potential is different from adding loop extrusion.

**Fig. S8.**
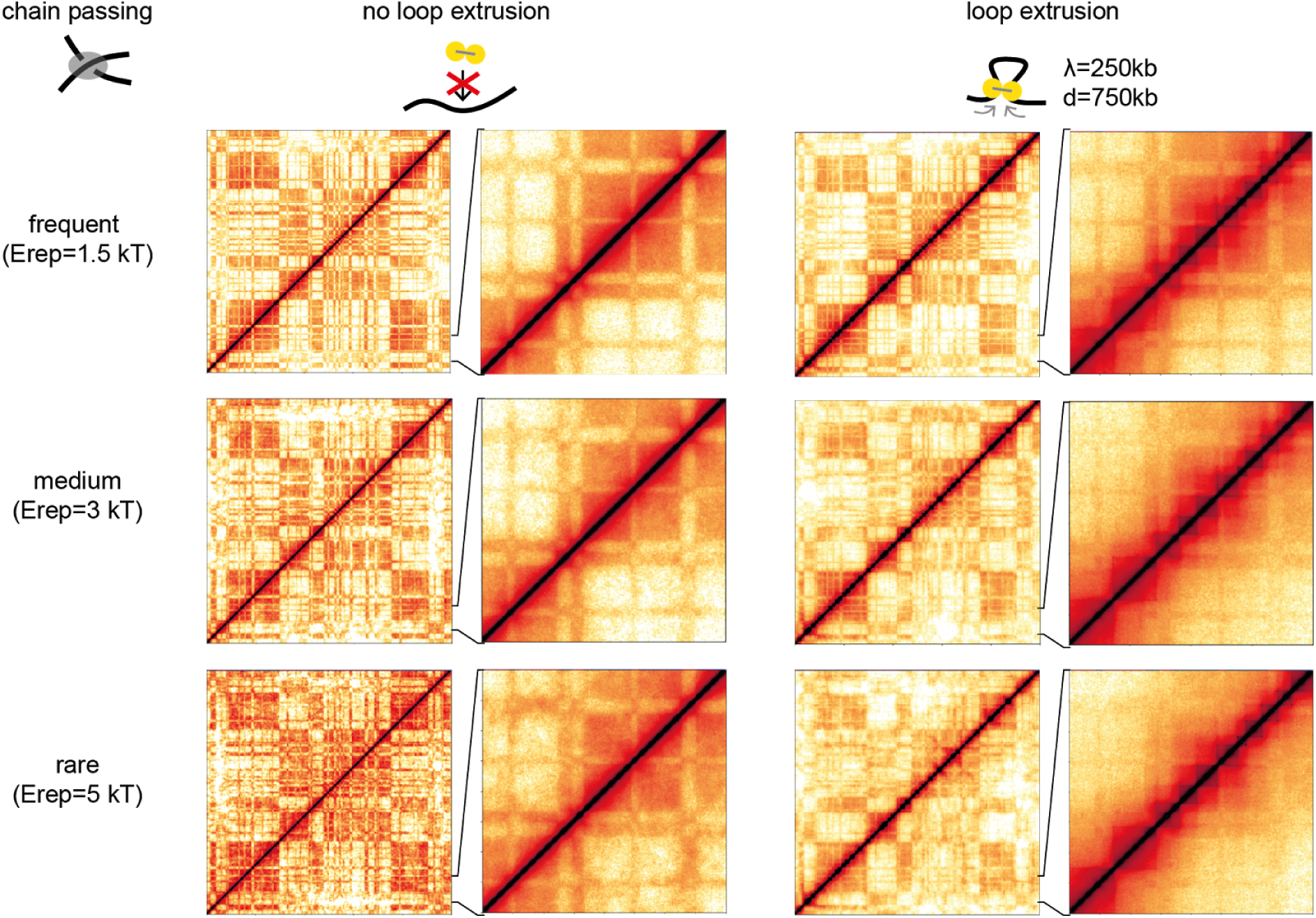
Hi-C maps from simulations for different degrees of chain passing (parametrized by E_rep_, the repulsive part of the monomer interaction potential), with and without loop extrusion.

**Fig. S9.**
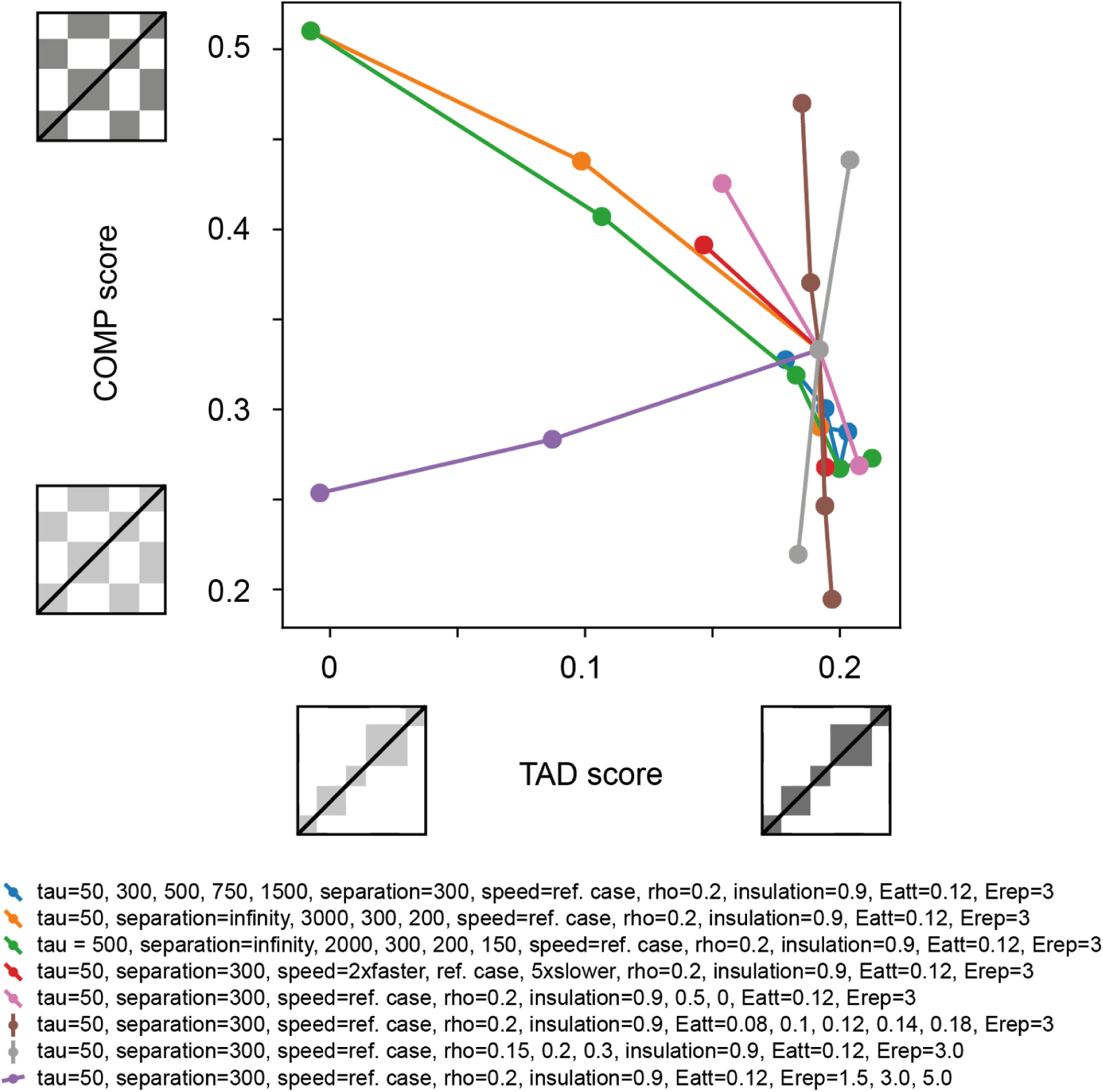
Same data as Fig. 6 in the main text, but with complete list of simulation parameters.

**Fig. S10.**
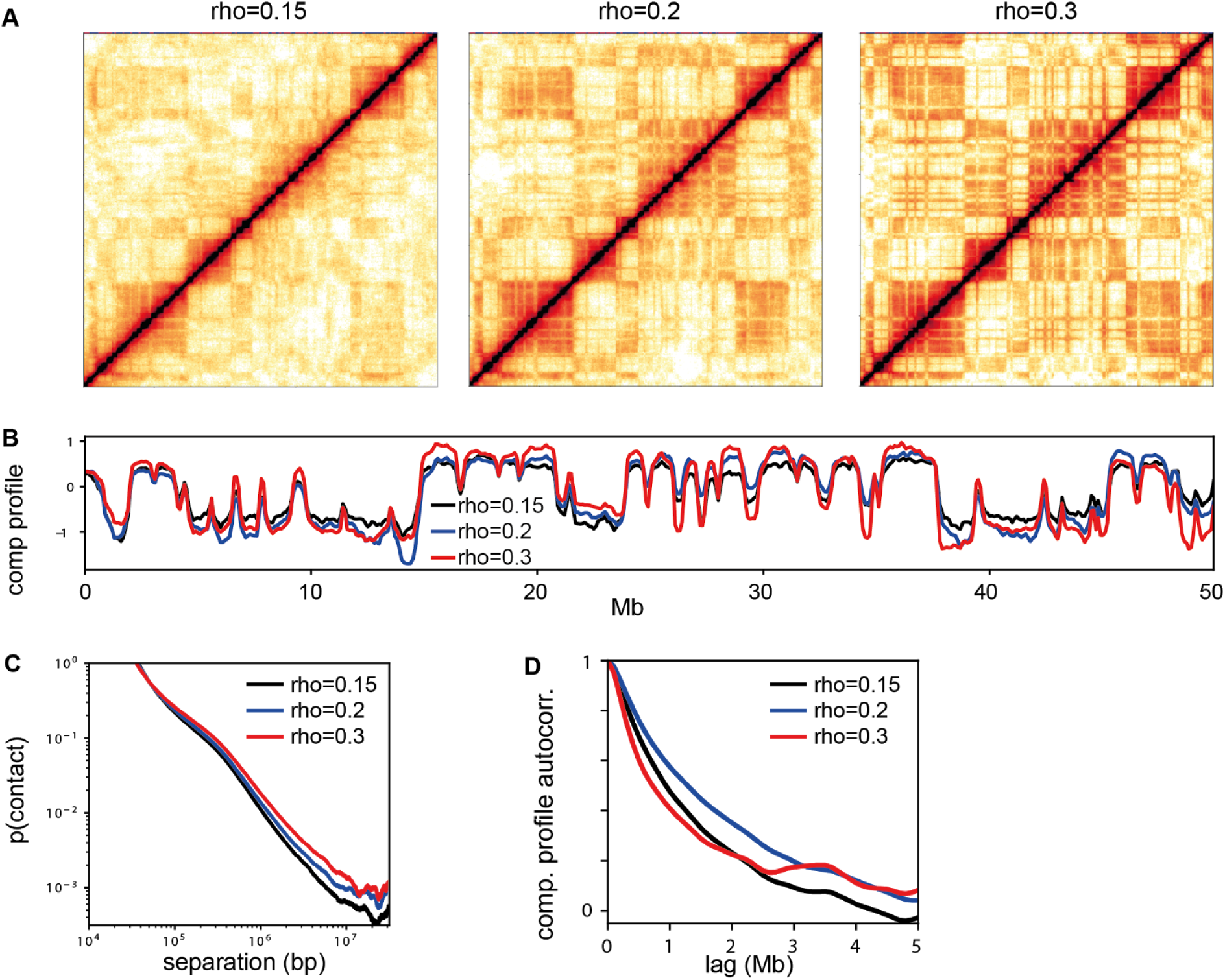
Contact matrices, compartment profiles, contact probability scaling, and compartment profile autocorrelation for different volume densities of our simulated chromatin fiber. All other parameters are as in our reference case in the main text. Changes in density are intended to reflect corresponding changes in nuclear volume (note that our simulations of 50 Mb of chromatin are performed in periodic boundary conditions).

